# Chemoenzymatic synthesis of sialylated and fucosylated mucin analogs reveals glycan-dependent effects on protein conformation and degradation

**DOI:** 10.1101/2025.04.15.648972

**Authors:** Amanda M. Wood, Casia L. Wardzala, Jessica R. Kramer

## Abstract

Mucin proteins are essential for life but are challenging to study due to their complex glycosylation patterns. Synthetic mimics have become vital tools for understanding and modulating the roles of mucins in human health and disease. We developed a chemoenzymatic approach to prepare polypeptide-based synthetic mucins displaying a variety of glycans with native linkages and orientations. By combining the polymerization of glycosylated amino acid *N*-carboxyanhydrides with enzymatic sialylation and fucosylation, we produced a tunable panel of synthetic mucins. These polymers were recognized by natural glycan-binding and glycan-degrading enzymes, providing insights into the structural preferences of these proteins. Glycan- and linkage-dependent effects on proteolysis were observed. Further, investigation of the influence of glycans on peptide backbone secondary structure revealed that both sialylation and linkage at Ser vs. Thr have profound effects on hierarchical conformation. Overall, our methodology offers versatile tools for exploring the diverse glycobiology of mucins.

## Introduction

Mucins are a family of glycoproteins that coat every wet surface in the human body and are foundational to mucus, tears, saliva, saliva, and the epithelial and endothelial glycocalyces.^1,2^ More than twenty human mucins have been identified with differing polypeptide sequences, but all characteristically feature a densely *O*-glycosylated domain rich in Pro, Thr, and Ser (PTS domains).^3–5^ Mucin glycosylation initiates with α-*N*-acetylgalactosamine (αGalNAc) attached to Ser or Thr. Additional sugars can be appended to the 2-, 3-, 4-, and/or 6-hydroxyls, with glycan chains frequently terminating in fucose (Fuc) or sialic acid (Sia).^6–8^ Due to their terminal location and multivalent display, Sia and Fuc are primed for diverse binding interactions. For example, these sugar groups play essential roles in regulation of extravasation, immune function, cancer, pathogen defense and interaction with symbiotic microbes.^9–11^ Yet, probing molecular details of these interactions has been challenging due to mucins’ inherent heterogeneity. Here, we present tunable and well-defined synthetic mucin analogs that display Sia and Fuc in their native chemical forms.

Biosynthesis of glycan-mature mucins takes place in the Golgi via glycosyltransferase enzymes.^12^ Eight core mucin di- and tri-saccharide structures have been identified, featuring αGalNAc linked with various regio- and stereo-chemistries to N-acetylglucosamine (GlcNAc), GalNAc, and/or galactose (Gal). These structures can be further diversified and commonly terminate with Sia and Fuc moieties, again linked with unique regio- and stereo-chemistries. The result is an impressive and densely packed array of chemical complexity.

These discrete mucin glycan structures, each possessing differing molecular shapes and chemical properties, confer unique biology as they vary with species, tissue, and disease. For example, distinct human tissue distributions of Sia α2,3- and α2,6-linked to Gal or GalNAc appear to contribute differently to viral infection, cancer progression, and receptor binding on both adaptive and innate immune cells.^13–20^ Similarly, Fuc α1,2-, α1,3-, and α1,4-linked to Gal or GlcNAc play differing roles in human blood group antigens, cancer, chronic respiratory disorders, and coronavirus infection.^21–25^

Due to the complexity of glycan chemistry and biology, glycan-dependent effects are typically correlative rather causative. Detailed molecular studies have been roadblocked by the intricacies of protein glycosylation biosynthesis which integrates diet, environment, and genetics into hundreds or even thousands of enzymes.^26–28^ Advances in genetic engineering have made strides in overcoming the challenges in expressing the large, highly repetitive protein sequences characteristic of mucins.^29,30^ Recent work tackled glycoengineering of mucins in a model human cell line and remarkable control over simple glycan patterning was reported.^31^ Despite these advances, precise control over glycan density and pattern, particularly with respect to regio- and stereo-specific sialylation and fucosylation, remains problematic. Characterization of glycan patterns in biologically-derived mucins is a challenge in and of itself, and scalability of niche recombinant materials has limitations. Synthetic mimics are poised to address these challenges and serve as tools to probe mucin biology.

A variety of mucin surrogates have been explored including short peptides, polysaccharides, and synthetic polymers.^32,33^ Polymeric mimics are particularly promising tools for further development since they can capture the glycan multivalency long-known to play key roles in diverse glycan binding events.^34–36^ However, these prior surrogates are generally poor substitutes due to their inability to achieve the molecular weights, multivalency, conformations, chemical linkages, or specific glycans found in nature.

To address the need for improved mucin surrogates, we have been developing peptide polymers that feature natural chemical linkages and adopt native protein secondary structures.^37–43^ These tunable synthetic mucins (synMUCs) are derived from α-amino acid *N*-carboxyanhydride (NCA) polymerization, which is a rapid and scalable (mg–kg) technique used in research, commodities, and pharmaceuticals.^44–47^ With application of modern polymerization initiators, polypeptides from ca. 10–1000 residues and with low dispersity (Ð<1.2) can be reproducibly prepared. Considering the importance of terminal Sia and Fuc glycans in mammalian biology, we sought to develop synMUCs that display these glycans with their various native linkage orientations and with tunable density.

Efforts towards multivalent Sia^48–52^ and Fuc^53–55^ display have focused on non-native backbones such as methacrylate, norbornene, methylvinylketone, lactic-co-glycolic acid. Fuc- and Sia-modified short peptides have also been described.^56–59^ As previously described, such surrogates have inherent limitations based on use of non-natural chemical groups or low molecular weight, thus affecting biological interactions from glycan orientation to lack of multivalency. For example, solid-phase-synthesis-derived peptides typically have only ca. 10 residues and only one or two glycosylation sites. Natural mucins are hundreds, or even thousands, of residues long and can display glycans at 50% or more of these sites.^3–5^ Similarly, natural mucin proteins take on ordered secondary structures that flexible synthetic polymers cannot adopt, surely affecting glycan display. We could find only a single example of α2,6 Sia display from a polypeptide backbone^60^ and could find no such reported systems for Fuc.

To achieve our goal of Sia- and Fuc-presenting synMUCs, we utilized NCA chemistry to prepare polypeptides bearing simple glycans which were substrates for regio- and stereo-selective sialylation and fucosylation via an efficient one-pot multi enzyme (OPME) strategy. After optimization of this chemoenzymatic strategy, we validated synMUC binding to native targets, probed glycan-dependent effects on peptide conformation, and investigated their proteolytic susceptibility to mucinase enzymes.

Characterization of glycan-dependent effects on conformation, binding, and degradation are highly relevant to human health and disease. For example, glycosylation changes associated with cancer, chronic respiratory diseases, or infections,^61–64^ could affect protein structure and have undiscovered functional outcomes in these pathologies. Similarly, mucinases, identified in diverse mucus-resident pathogenic and commensal microbes,^65–71^ impact mucin-microbe relationships and offer therapeutic opportunities. There has been a recent explosion of interest in mucinases for application as therapeutics and analytical tools.^72–76^ Here, we report glycan-dependent effects on glycopeptide conformation and mucinase degradation which may shed light on the structure and function of natural mucin glycoproteins and inform on the broader application of mucinases in research and therapeutics.

## Results and Discussion

### Chemical synthesis of synMUCs with simple glycosylation

For preparation of the synMUC backbones, we chose NCA polymerization since the resulting materials are composed entirely of amino acids and sugars with their natural linkages, and native protein secondary structures can be adopted. ^37–39^ The NCA method also allows precise tuning of molecular weight (MW), and peptide composition including glycan density and glycan identity. For this study, we selected Ser/Thr conjugates of αGalNAc, βGal, and β1,3Gal-αGalNAc (core 1) (Figure 2A). These glycans are abundant in mucins, both as standalone moieties and components of core structures, and are substrates for enzymatic sialylation and fucosylation. Further, these glycans are relevant to human disease. For example, in carcinoma tissues the Thomsen-nouveau (Tn) antigen, αGalNAc, the Thomsen-Friedenreich (Tf) antigen, β1,3Gal-αGalNAc, and sialyl-Tn appear to play roles in tumorigenesis, immune evasion, and more.^77–79^

**Figure 1:**
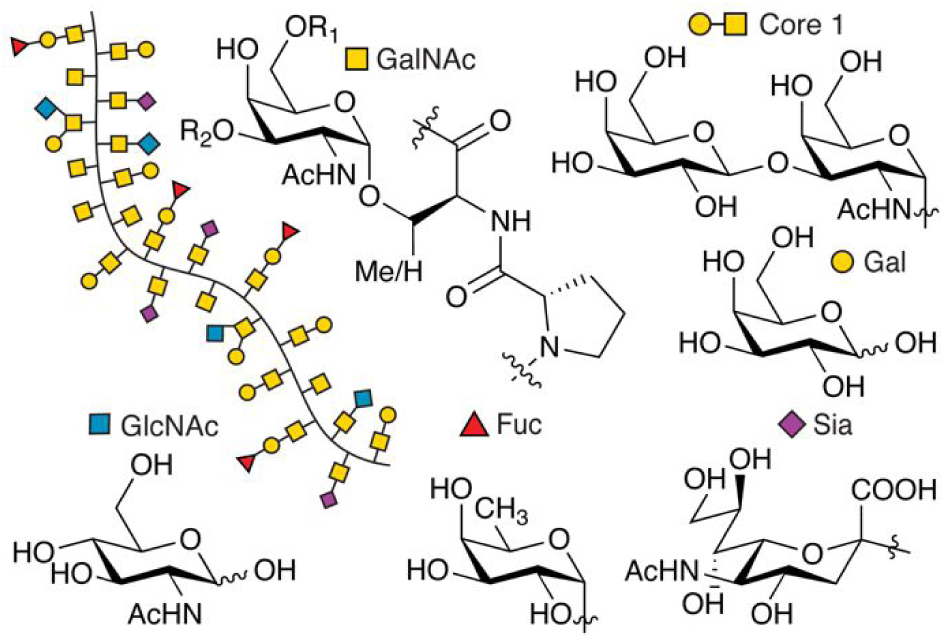
Structure of proline-, threonine-, and serine-rich mucin PTS domains with *O*-linked α-*N*-acetylgalactosamine, which can be elaborated with subsequent glycans such as galactose, *N*-acetylglucosamine, or terminal fucose and sialic acids.

**Figure 2:**
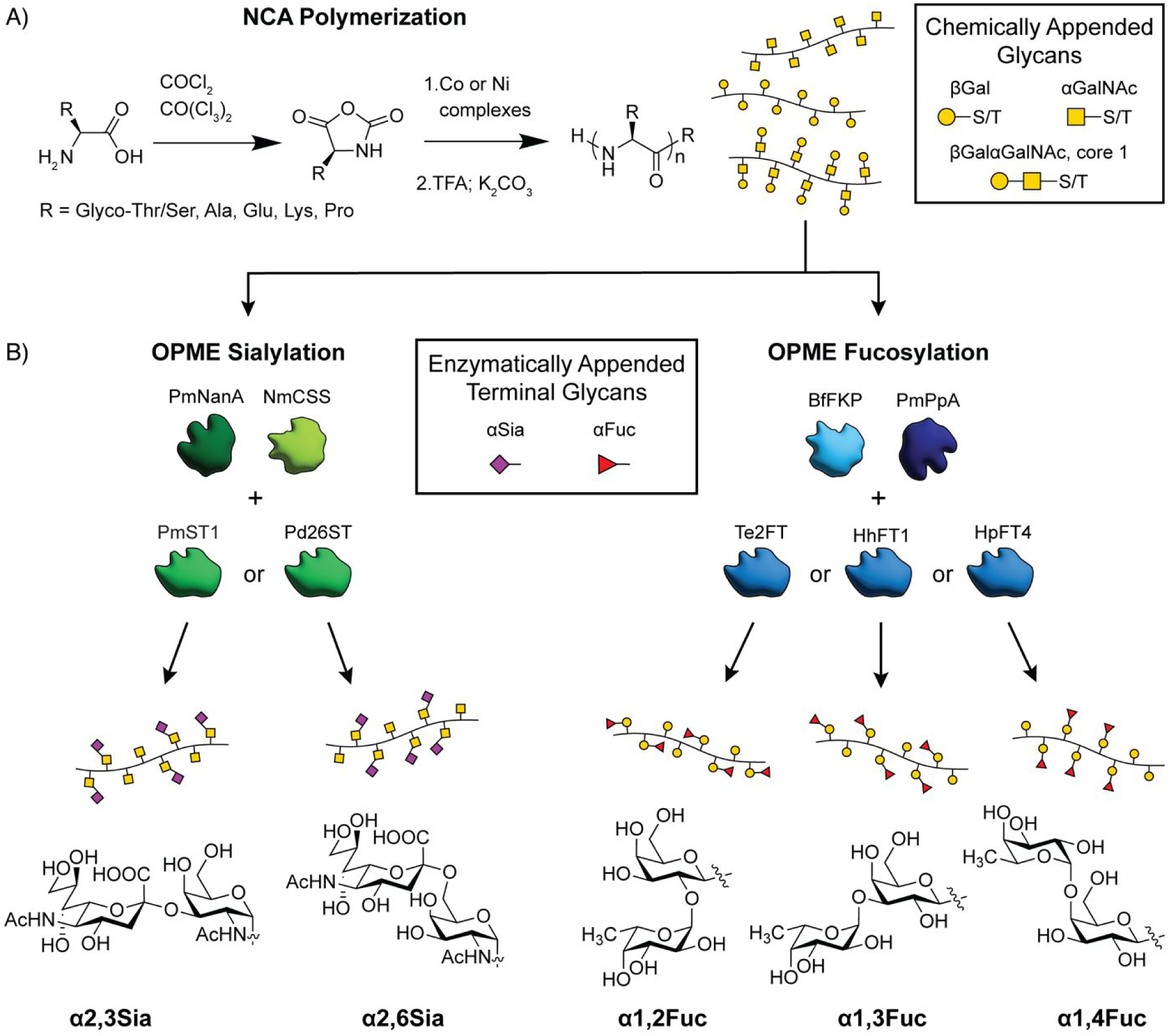
Chemoenzymatic preparation of synthetic mucins (synMUCs) with regio- and stereo-specific presentation of mucin glycans. A) Chemical synthesis of synMUC glycopolypeptide backbones by polymerization of amino acid NCAs, with readily tunable chain length, glycan density, and amino acid composition. B) One-pot multi-enzyme (OPME) reactions append terminal α2,3- or α2,6-linked sialic acid or α1,2-, α1,3-or α1,4-linked fucose onto synMUC backbones.

Established methodology was utilized to prepare the Ser and Thr glyco-conjugates and convert them to NCAs (Figure 2A).^37–43^ To initiate NCA polymerization, we employed nickel or cobalt based organometallic complexes known to offer fast kinetics, side reaction suppression, controlled and living polymerization, and to yield high molecular weight (MW) polymers of low dispersity.^47,80^ The nickel complexes also allow for installation of clickable groups at the initiation site^38,42^, which we used later for attachment to 96-well plates for binding assays. With both initiators, the peptide termini are amine groups primed for conjugation to biotin, fluorophores, or other molecules of interest.

We hypothesized that glycan density and proximity to charged groups or sterically restricted residues could affect activity of mucin-modifying glycosyltransferases and mucin-degrading glycoproteases. To address these questions, we prepared synMUC variants with differing backbone structures. βGal-, αGalNAc-, and β1,3Gal-αGalNAcSer/Thr NCAs were singularly polymerized to yield homopolypeptides with 100% glycosylation. We also mixed the glyco-NCAs in various ratios with “spacer” amino acid NCAs to prepare polypeptides with varying densities of glycosylation, differing ionic charges, and different steric complexity (Table 1).

**Table 1:** Representative data for preparation of polypeptide-based synMUCs by NCA polymerization. Polymerizations were conducted under inert atmosphere in THF and initiated with [a] (PMe_3_)_4_Co or [b] Ni azido-amidoamidate catalyst when clickable termini were desired. Polymerizations generally proceeded at ambient temperature; however specific cases (*) were aided by heating at 50 °C. [c] Abbreviated structure name. Observed [d] number average molecular weight, M_n_, [e] degree of polymerization (DP), and [f] dispersity, Đ, as determined by SEC/MALS/RI in 0.1M LiBr in DMF.

| SynMUC | Abbrv.<sup>[c]</sup> | $M_n$<sup>[d]</sup> | DP<sup>[e]</sup> | $\bar{D}$<sup>[f]</sup> |
| --- | --- | --- | --- | --- |
| $(\text{Gal}\beta\text{Ser})_n$<sup>[a]</sup> | 100% $\beta\text{Gal}^S$ | 45,070 | 108 | 1.34 |
| $(\text{Gal}\beta\text{Thr})_n$<sup>[a]</sup> | 100% $\beta\text{Gal}^T$ | 43,400 | 101 | 1.18 |
| $(\text{Gal}\beta\text{Ser}_{0.5}\text{-s-Glu}_{0.25}\text{-s-Ala}_{0.25})_n$<sup>[a]</sup> | 50% $\beta\text{Gal}^{\text{SEA}}$ | 32,760 | 120 | 1.04 |
| $(\text{Gal}\beta\text{Thr}_{0.5}\text{-s-Glu}_{0.25}\text{-s-Ala}_{0.25})_n$<sup>[b]</sup> | 50% $\beta\text{Gal}^{\text{TEA}}$ | 30,590 | 112 | 1.17 |
| $(\text{Gal}\beta\text{Thr}_{0.5}\text{-s-K}_{0.25}\text{-s-Ala}_{0.25})_n$<sup>[a]</sup> | 50% $\beta\text{Gal}^{\text{TKA}}$ | 31,940 | 110 | 1.33 |
| $(\text{Gal}\beta\text{Thr}_{0.25}\text{-s-Glu}_{0.38}\text{-s-Ala}_{0.38})_n^{[b]}$ | 25% $\beta\text{Gal}^{\text{TEA}}$ | 22,800 | 112 | 1.03 |
| $(\text{Gal}\beta\text{Thr}_{0.25}\text{-s-Glu}_{0.25}\text{-s-Ala}_{0.25}\text{-s-Pro}_{0.25})_n^{[b]*}$ | 25% $\beta\text{Gal}^{\text{TEAP}}$ | 20,880 | 106 | 1.12 |
| $(\text{GalNAc}\alpha\text{Ser})_n^{[a]}$ | 100% $\alpha\text{GalNAc}^{\text{S}}$ | 42,690 | 103 | 1.30 |
| $(\text{GalNAc}\alpha\text{Thr})_n^{[a]}$ | 100% $\alpha\text{GalNAc}^{\text{T}}$ | 43,550 | 101 | 1.30 |
| $(\text{GalNAc}\alpha\text{Ser}_{0.5}\text{-s-Glu}_{0.25}\text{-s-Ala}_{0.25})_n^{[a]}$ | 50% $\alpha\text{GalNAc}^{\text{SEA}}$ | 26,650 | 98 | 1.18 |
| $(\text{GalNAc}\alpha\text{Ser}_{0.5}\text{-s-K}_{0.25}\text{-s-Ala}_{0.25})_n^{[a]}$ | 50% $\alpha\text{GalNAc}^{\text{SKA}}$ | 26,070 | 92 | 1.32 |
| $(\text{GalNAc}\alpha\text{Ser}_{0.25}\text{-s-Glu}_{0.38}\text{-s-Ala}_{0.38})_n^{[a]}$ | 25% $\alpha\text{GalNAc}^{\text{SEA}}$ | 19,400 | 115 | 1.24 |
| $(\text{GalNAc}\alpha\text{Ser}_{0.25}\text{-s-Glu}_{0.25}\text{-s-Ala}_{0.25}\text{-s-Pro}_{0.25})_n^{[b]*}$ | 25% $\alpha\text{GalNAc}^{\text{SEAP}}$ | 19,970 | 104 | 1.06 |
| $(\text{Gal}\beta\text{-GalNAc}\alpha\text{Thr}_{0.25}\text{-s-Glu}_{0.38}\text{-s-Ala}_{0.38})_n^{[b]*}$ | 25% $\beta\text{Gal}\text{-}\alpha\text{GalNAc}^{\text{TEA}}$ | 27,290 | 99 | 1.53 |

For spacer residues, we chose Ala, Glu, Lys, and Pro which naturally occur in mucin PTS domains. Ala NCA was selected due to its small, neutral, and non-reactive side-chain, Glu to achieve anionic charge, Lys to achieve cationic charge, and Pro for its highly restrictive φ and ψ angles and abundance in native mucin glycodomains. For consistent comparison of glycopolypeptides with varied compositions, we prepared all structures with chain lengths of around 100 residues. Polymerizations were monitored by attenuated total reflectance Fourier-transformed infrared spectroscopy (ATR-FTIR) and characterized by size exclusion chromatography coupled with multi-angle light scattering and refractive index detectors (SEC/MALS/RI). As expected from previously established methodology^37–43^, polymerizations were high yielding and gave low dispersity materials (See SI for representative analytical data).

### Enzymatic modification of synMUCs with Sia and Fuc glycans

Extensive literature has associated α2,3- and α2,6-linked Sia (*N*-acetylneuraminic acid, Neu5Ac) groups with cancer, immunity, and human and zoonotic infections.^13–16,78,81^ Therefore, we selected these structures to append via native linkages to αGalNAc (Figure 2B). Similarly, overexpression of Fuc-bearing glycans has been correlated to asthma, cystic fibrosis, aggressive pancreatic and lung cancers, and altered microbial interactions.^10,82–91^. Hence, we targeted installation of native α1,2-, α1,3-, and α1,4-Fuc linkages to βGal (Figure 2B). Considering the complexity of selectively installing Sia and Fuc regio- and stereo-selectively via chemical synthesis, we investigated enzymatic glycosylation of our NCA-derived mono- and di-saccharide bearing synMUCs.

Identified in both mammals and bacteria, sialyltransferases (STs) and fucosyltransferases (FTs) utilize cytidine 5′-monophosphate-Sia (CMP-Sia) or guanosine 5′-diphosphate-Fuc (GDP-Fuc), respectively, to sialylate or fucosylate glycan substrates. We first investigated the in vitro reaction of our synMUCs with commercially available sialyltransferases (Photobacterium damsela α-2,6-sialyltransferase and Pasteurella multocida α-2,3-Sialyltransferase from Sigma) and CMP-Sia (Nacalai USA). However, we found the reactions to be very low yielding since the activated sugar has poor shelf stability due to hydrolysis and some commercial enzymes were of poor quality and could not be dissolved in water. To address such issues, optimize yields, and improve economic viability of enzymatic glycosylations, Chen and coworkers pioneered one-pot multi-enzyme (OPME) glycosylation reactions that generate the activated CMP-Sia or GDP-Fuc donors in situ from economical glycan precursors.^22,23,92–99^ In some cases, they report remarkable gram-scale production. Inspired by their work, we sought to expand their OPME methodology from free glycans to polypeptide-bound glycan substrates.

In the OPME sialylations, *N*-acetyl mannosamine (ManNAc) is first converted to Sia by sialic acid aldolase (PmNanA), followed by conversion to CMP-Sia by CMP-sialic acid synthetase (NmCSS) in the presence of cytidine 5′-triphosphate (CTP). In fucosylation reactions, GDP-Fuc is formed from L-fucose, adenosine 5′-triphosphate (ATP), and guanosine 5′-triphosphate (GTP) using bifunctional enzyme (BfFKP) that has both L-fucokinase and GDP-Fuc pyrophosphorylase activities. Inorganic pyrophosphatase (PmPpA) is also present to drive the reaction forward. Necessary in each case is a relevant FT or ST enzyme. For STs, we selected proteins from *Pasteurella multocida* and *Photobacterium damsela* since expression has been previously reported in *E. coli* and their α2,3- and α2,6-linkage specificities have been established.^97,100^ Similarly, we selected FTs originating from *Thermosynechococcus elongatus*, *Helicobacter hepaticus*, and *Helicobacter pylori* for their ability to install α1,2-, α1,3-, or α1,4-Fuc, respectively.^23,96,101^ We recombinantly expressed the set of nine enzymes on the mg scale using modified literature protocols (Table 2). All enzymes were 6x-His-tagged to allow for purification via Ni^2+^ or Co^2+^ affinity columns (see SI).

**Table 2.** List of enzymes utilized for OPME sialylation or fucosylation of synthetic mucins at Gal or GalNAc.

| Enzyme | Origin Species | Abbrev. | Glycan | Yield (mg/L) |
| --- | --- | --- | --- | --- |
| bifunctional L-fucokinase, GDP-fucose pyrophosphorylase | *Bacteroides fragilis* | BfFKP | - | 24.0 |
| Inorganic pyrophosphatase | *Pasteurella multocida* | PmPpA | - | 183.0 |
| α1,2-fucosyltransferase | *Thermosynechococcus elongatus* | Te2FT | α1,2-Fuc | 7.4 |
| α1,3-fucosyltransferase | *Helicobacter hepaticus* | HhFT1 | α1,3-Fuc | 27.0 |
| α1,4-fucosyltransferase | *Helicobacter pylori* | HpFT4 | α1,4-Fuc | 3.4 |
| sialic acid aldolase | *Pasteurella multocida* | PmNanA | - | 352.1 |
| CMP-sialic acid synthetase | *Neisseria meningitidis* | NmCSS | - | 411.7 |
| α2,3-sialyltransferase | *Pasteurella multocida* | PmST1 | α2,3-Sia | 226.8 |
| α2,6-sialyltransferase | *Photobacterium damsela* | Pd2,6ST | α2,6-Sia | 23.9 |

With purified enzymes in hand, OPME sialylation and fucosylation of chemically synthesized synMUCs was examined. Considering the large number of possible combinations, we did not attempt enzymatic glycosylation of each of the 14 synMUC compositions with each of the 5 enzymes. Instead, we chose representative samples for comparison. Detailed optimized conditions for the OPME reactions are reported in the SI. In brief, we used 1.5 equivs of monosaccharide donor and 1.3–1.5 equivs of NTP per mole of acceptor glycan. For fucosylations, we used 0.067 mg/mL PmPpA and 0.1 mg/mL BfFKP with 0.1 mg/mL Te2FT, HhFT1, or HpFT4. For sialylations, we used 0.1 mg/mL NmCSS and 0.2 mg/mL PmNanA with 0.2 mg/mL PmST1 or Pd2,6ST. In all cases, reactions were in Tris buffer with MgCl_2_. For all synMUC glycosylation reactions, 6xHis tagged enzymes were removed from the reaction mixture via incubation with Ni-NTA magnetic beads. SynMUCs were purified by dialysis against ultrapure water and lyophilized to yield white powders.

Key to characterizing the synMUCs was an accurate method of glycan analysis. Therefore, we undertook multiple approaches to tackle this non-trivial task. Analysis by ATR-FTIR and NMR confirmed the presence of the correct functional groups and we could see appropriate increases in the glycan integrals (see SI). However, the overlap of glycan and amino acid spectral signatures and broad peaks due to the polymeric nature of the materials convoluted accurate calculation of the extent of fucosylation and sialylation from peak integrations. We also explored commercially available glycan analysis kits for Sia and Fuc (Sigma, Megazyme). The kits were useful to detect relative changes between reactions with differing compositions. However, the results were qualitative as Sia assays consistently reported higher than theoretical yields and the Fuc assay kit suffered from specificity issues where control samples containing only Gal yielded results positive for Fuc.

To more accurately confirm the predicted structures and to quantify the amounts of added carbohydrates, we turned to gas chromatography-mass spectrometry (GC-MS), high-performance anion exchange chromatography (HPAEC), X-ray photoelectron spectrometry (XPS), and SEC/MALS/RI. These techniques are attractive since only µg quantities are required.

For GC-MS, amino and neutral sugars were analyzed after treatment of synMUCs with acidic methanolysis and conversion to methyl glycosides followed by trimethylsilyl (TMS) derivatization. Since Sia degradation can occur during acidic methanolysis, detection of this glycan was attempted using HPAEC. To avoid interference of amino acids on HPAEC, synMUCs were treated with an α-2-3,6,8,9-neuraminidase A to release Sia residues from the polypeptide prior to analysis. MS results confirmed that Sia and Fuc groups had indeed been enzymatically appended to our polymers (see SI). However, we could not accurately determine Sia and Fuc content using this method. Glycan ratios were inconsistent for our core 1 synMUC samples where 1:1 ratios of Gal:GalNAc were expected and glycan yields were consistently lower than observed by XPS, SEC, NMR and the commercial assays. We suspect this is due to variability in efficiency of methanolysis, silylation, or sialidase cleavage.

XPS was an attractive alternative since the technique can use intact glycopolypeptides with no chemical reactions needed prior to analysis, and very small sample mass is required. Intact synMUCs were analyzed for elemental composition and high-resolution scans were deconvoluted into characteristic peaks for various bond species. The results of these analyses and representative spectra are shown in Figure 3. Additional data can be found in the SI. Appending of Fuc or Sia residues generates changes in the number, and consequently ratios, of C–C, C═O, C–O, and C–N bonds. Therefore, we compared the spectra of the synMUCs before and after the enzymatic reactions and calculated the fold-change in these bonds. C 1s curves were deconvoluted into their constituent C-C, C-N/C-O, and C=O peaks and plotted under their calculated C 1s envelope curve. O-C=O peaks were added when apparent. To correct for adventitious carbon contamination, C=O peaks were normalized between samples when appropriate. Glycosylation efficiency was defined as the observed number of Sia or Fuc groups as compared to the total number of potential Gal or GalNAc acceptors, and was calculated via the observed fold change in C-N/C-O / C=O vs. the theoretical maximum.

**Figure 3.**
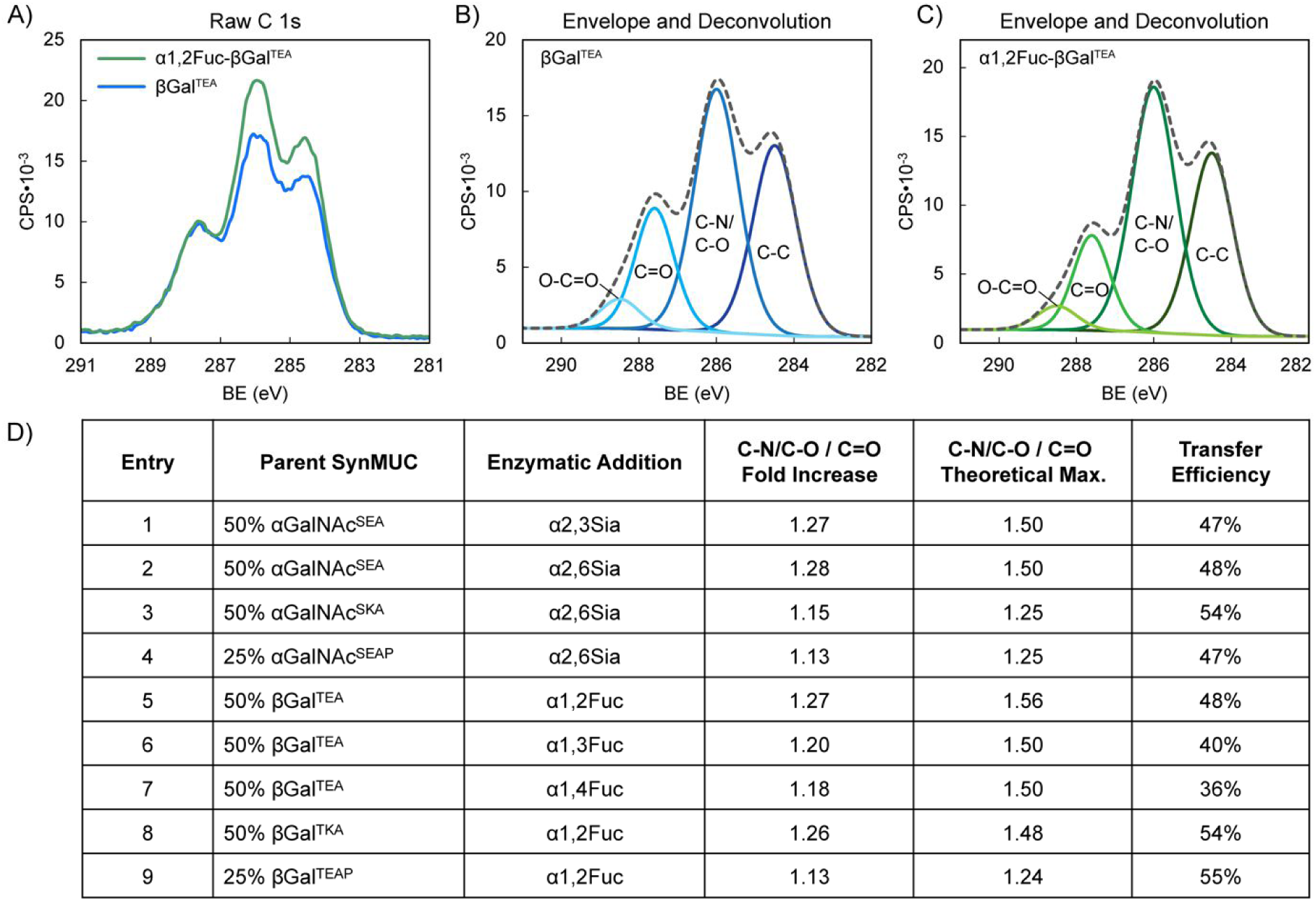
Application of XPS to quantify the extent of enzymatic synMUC fucosylation or sialylation via changes in bond species ratios. Representative XPS raw C 1s spectra of A) 50% βGal^TEA^ before and after α1,2 fucosylation, B−C) Deconvolution of C 1s curves into their constituent C-C, C-N/C-O, and C=O peaks plotted under their calculated C 1s envelope curve. D) Summary of XPS results for enzymatically sialylated and fucosylated synMUCs.

To further confirm the accuracy of the XPS data, we selected a subset of samples to analyze by aqueous SEC/MALS/RI before and after the OPME reactions. For this technique, we expected to see a molecular weight shift due to the extra mass of the Fuc or Sia residues. Number average molecular weight (M_n_) and dispersity were calculated from the MALS and RI data using Astra 7 software. Figure 4 reports the results of these analyses with representative chromatograms included. Due to similar interactions of the glycopolypeptides with the column stationary phase, major shifts in elution time are not expected and elution time is not a factor in the calculations. We were pleased to see unimodal glycopolypeptide distributions after the OPME reactions, indicating chains remain intact and only one species is present. Gratifyingly, side by side comparison of the SEC and XPS data revealed the two methodologies are in relatively close agreement.

**Figure 4:**
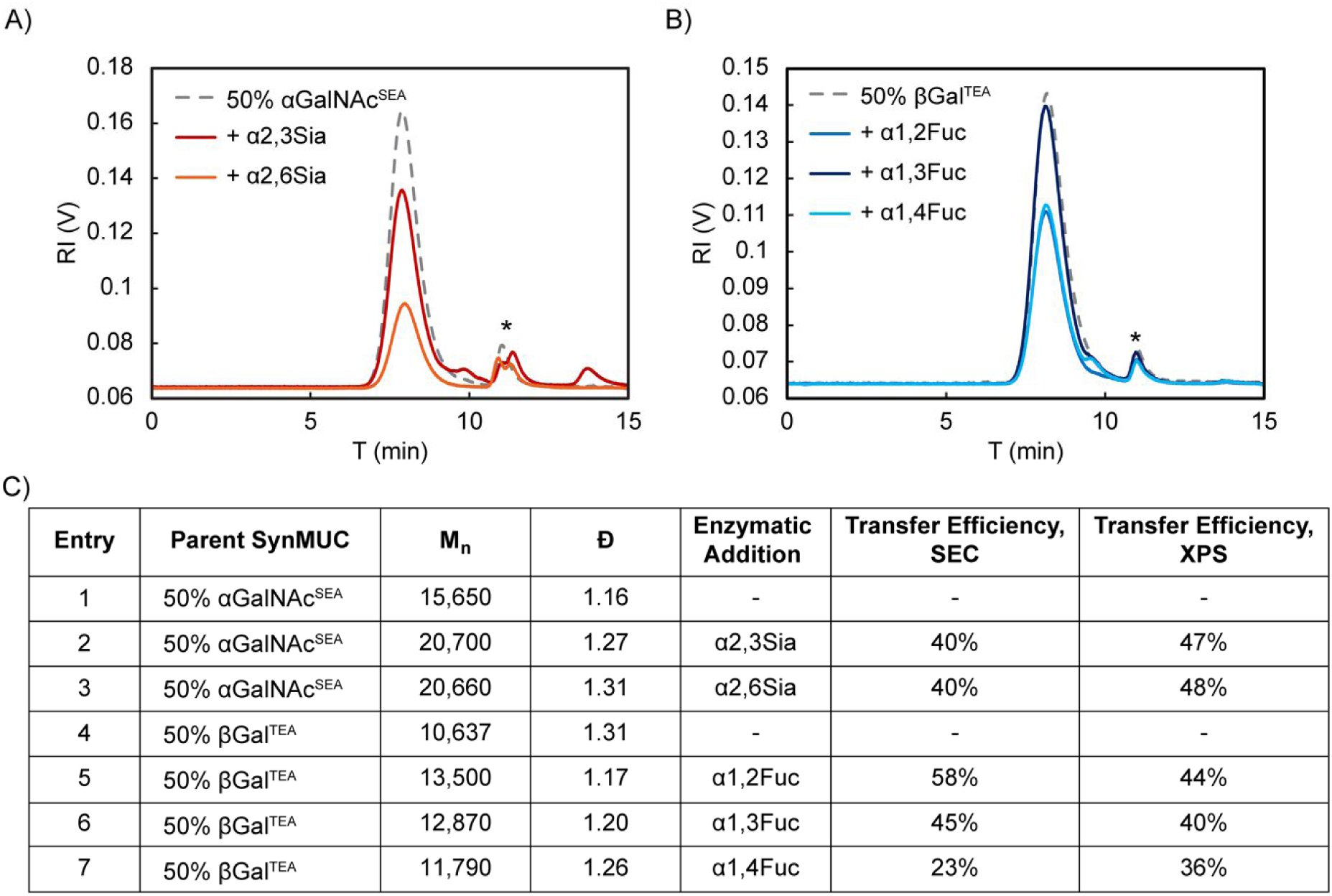
Analysis of glycopolypeptide molecular weights by SEC/MALS/RI and comparison to XPS. A) RI traces of the parent 50% αGalNAc^SEA^ as compared to α2,3- and α2,6-sialylated products. B) RI traces of the parent 50% βGal^TEA^ as compared to α1,2-, α1,3-, and α1,4-fucosylated products. In A, B) *Denotes solvent peak.C) Analytic results of the SEC/MALS/RI experiment as compared to XPS.

For the identical 50% glycosylated anionic αGalNAc^SEA^ acceptor polypeptide, efficiency of Sia transfer was essentially identical for both the α2,3 and α2,6 linkages generated by PmST1 and Pd2,6ST (47% and 48%, respectively, by XPS and both 40% by SEC, Fig. 3D, entries 1, 2 and Fig. 4C, entries 2, 3). To probe the effect of neighboring residue charge or steric effect, we used Pd2,6ST to sialylate acceptor polypeptides where anionic Glu was fully replaced with cationic Lys or where 25% of the residues were replaced by Pro (Fig. 3B, entries 2, 3). Glycan transfer efficiency was unaffected by the addition of Pro (47%, Fig. 3D, entry 3), which is rational considering that mucins are naturally rich in Pro. Sialylation efficiency improved slightly for the Lys containing peptide (54%, Fig. 3D, entry 4). We speculate this could be due to direct or allosteric stabilization of the enzyme active site, or simply favorable electrostatic interaction since the ST enzyme has an isoelectric point of 4.88 (Sigma-Aldrich^102^). Our results align nicely with those of the aforementioned report of α2,6-sialylation a 50%-lactose-Ser/Ala copolypeptide by Bertozzi and coworkers.^60^ They used a two-enzyme system of Pd2,6ST and NmCSS in the presence of Sia and reported ca. 50% efficiency as determined by SEC and NMR. They also reported unsuccessful attempts at α2,3-sialylation with a variety of STs, including PmST1. In our hands, using the OPME system with in-house expressed PmST1, we achieved excellent α2,3 sialylation yields for our synMUCs.

Fucosylation of an identical 50% βGal^TEA^ acceptor glycopolypeptide by the enzymes Te2FT, HhFT1, and HpFT4, which yield α1,2-, α1,3-, and α1,4-linkages, respectively, were relatively consistent (36–44% by XPS, Fig. 3D, entries 5–7; 23–58% by SEC, Fig. 4C, entries 5–7). To again probe steric or charge effects, we used Te2FT to fucosylate similar Lys or Pro containing backbones (Fig. 3D, entries 8, 9). Proline appeared to increase the favorability of the acceptor peptide since efficiency of fucosylation 25% βGal^TEAP^ increased slightly as compared to 50% βGal^TEA^ (55% vs. 48%), though we acknowledge this could be due to the lower glycan density also reducing steric hinderance. Similar to the sialylation case, lysine modified 50% βGal^TKA^ was a slightly more favorable substrate for Te2FT than the Glu containing polymer with 54% vs. 48% of the sites fucosylated (Figure 3D, Entries 3, 8).

The combination of NCA polymerization with OPME methodology is robust for generation of a variety of glycopolypeptide compositions. We are pleased to present first-in-kind materials with regio- and stereo-specific multivalent display of sialyl-Tn and fucose from native polypeptide backbones. In native mucins, reported Sia and Fuc content, as quantified by NMR or gas or liquid chromatography coupled to MS, is variable depending upon the MUC type and source tissue.^103–106^ Representative values for Sia and Fuc content, reported as % of total monosaccharide composition, ranged from 6–24% for Sia and 5–30% for Fuc. However, as seen in our chemically-defined materials, these data can be affected by the differing abilities of glycan structures to derivatize, enzymatically cleave, or ionize. In any case, we expect that the Sia and Fuc levels in our synMUCs are biologically relevant. Overall, we conclude the OPME reactions open doors to functional and tunable mucin mimics with application in probing the diverse biology of Fuc and Sia.

### Evaluation of functional glycan binding

Considering our proposed application of synMUCs as surrogates for native mucins, we sought to evaluate functional binding of the presented Sia and Fuc glycans. For proof-of-concept experiments, we selected commercially available biotinylated lectin proteins wheat germ agglutinin (WGA) and Ulex Europaeus agglutinin I (UEA-I) since these have previously been applied to mucin studies^107,108^. UEA-I has specificity for fucosylated glycans^109–111^, while WGA is reported to have specificity for Sia^110–113^. For binding assays, we utilized a dot blot format where synMUCs or control reagents were spotted on nitrocellulose in triplicate at varied concentrations. Membranes were washed, blocked with bovine serum albumin (BSA), and then biotinylated lectin was added. After washing, dot blots were incubated with streptavidin-IR680 fluorophore, washed, and imaged at 700 nm. Blot images were analyzed using ImageJ and fluorescence intensity was quantified. A representative dot bot is shown in Fig. 5A and additional data can be found in the SI.

**Figure 5.**
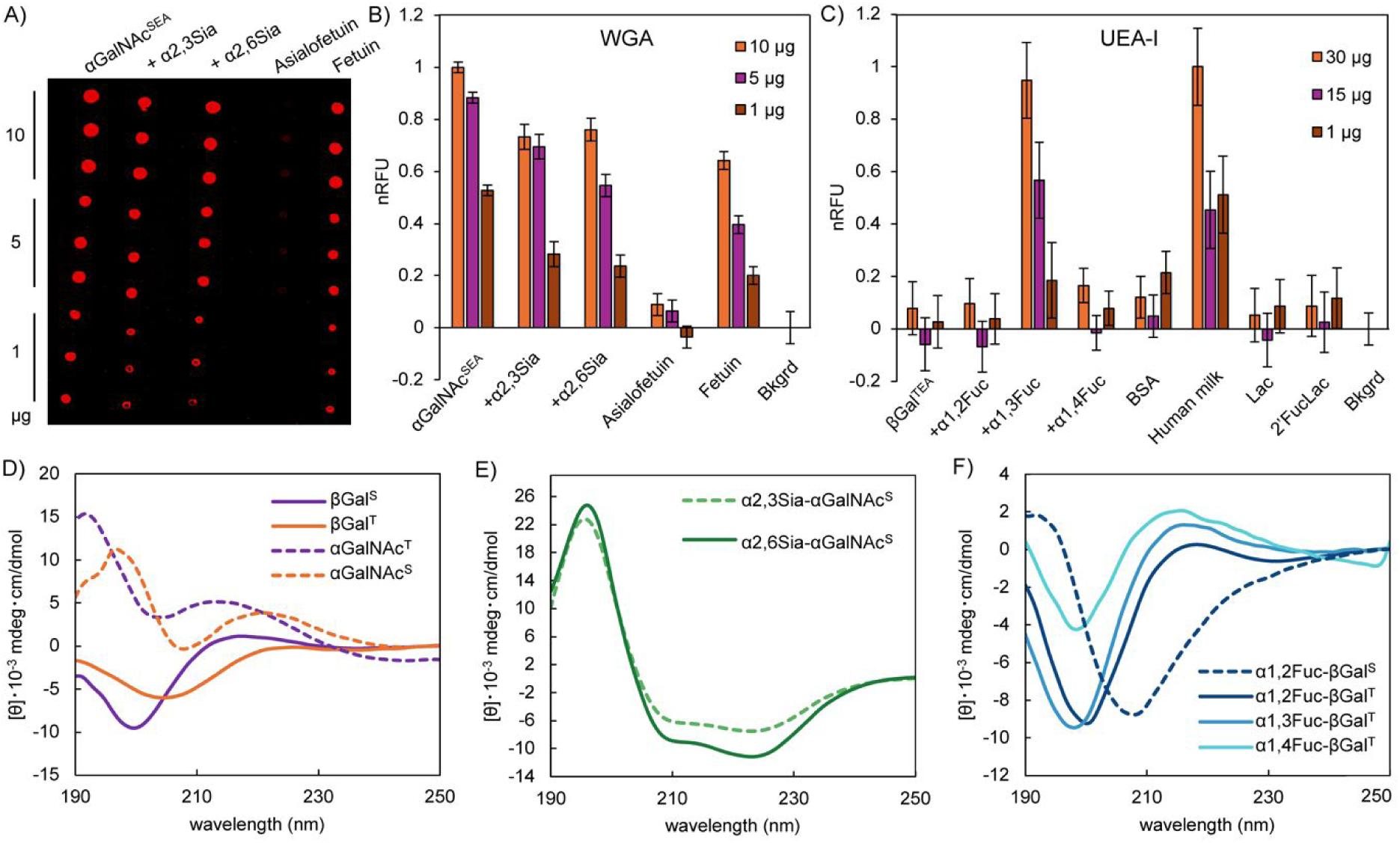
A) Representative lectin dot blot of parent synMUC αGalNAc^SEA^, enzymatically sialylated products, and controls. B) Quantified results of WGA lectin dot blot for 50% glycosylated αGalNAc^SEA^ parent synMUC, enzymatically sialylated products, and controls fetuin and asialofetuin. C) Quantified results of UEA-1 lectin dot blot for 50% glycosylated αGal^TEA^ parent synMUC, enzymatically fucosylated products, and controls BSA, human milk, Lac, and 2-fucosylLac. B, C) Error bars designate mean and standard deviation. D–F are circular dichroism spectra of 100% glycosylated polypeptides in Milli-Q water at 25 ° C and are reported in molar ellipticity. D) Comparison of the spectra of βGal vs. αGalNAc attached to Ser vs. Thr. E) Comparison of the spectra of various fucosylated regioisomers of βGal attached to Thr or Ser. F) Comparison of the spectra of sialylated regioisomers of αGalNAc^S^.

For the WGA dot blot, we examined binding to the 50% glycosylated αGalNAc^SEA^ parent synMUC as compared to the α2,3Sia-αGalNAc^SEA^ and α2,6Sia-αGalNAc^SEA^ enzymatic products (Fig. 5B). For controls, we chose commercially-available fetuin and asialofetuin and all samples were compared to membrane background signal. Signal from the controls was as expected, where binding was substantially higher for fetuin than asialofetuin. Signal above background for asialofetuin was expected, as WGA also has reported affinity for GlcNAc residues. We found that WGA had excellent binding to both the parent synMUC and the sialylated synMUCs. No preference for α2,3Sia vs α2,6Sia was observed and the lectin actually had a slightly stronger binding preference for the parent synMUC displaying only αGalNAc. This is not entirely unexpected considering literature reports of promiscuous binding to GlcNAc and GalNAc.^110,111^

Lectin dot blots with UEA-I utilized 50% glycosylated βGal^TEA^ as the parent synMUC, which was compared to the α1,2-, α1,3-, and α1,4-fucosylated products (Fig. 5C). For controls, we spotted BSA, which has no expected fucosylation; human milk, which is rich in diverse fucosylated glycans; lactose (Lac); and 2-fucosylLac (2’FucLac). For the controls, UEA-1 bound strongly to human milk as compared to BSA. Surprisingly, the lectin did not have any preference for 2’FucLac over Lac. We found that UEA-1 strongly preferred our α1,3Fuc-αGal^TEA^ polypeptide over the α1,2- and α1,4-Fuc regioisomers, or the parent βGal^TEA^ polymer.^109–111^ We found this result interesting because UEA-I is reported to prefer α1,2Fuc.^111,114^ However, we note prior lectin-glycan binding assays may not have directly compared these regioisomers or may have used free glycans not attached to a peptide where conformational effects could play a role.

Over the course of examining glycan-specific binding, we also attempted enzyme-linked lectin and immunosorbent assays (ELLA, ELISA) where azide-terminal glycopolypeptides were conjugated to alkyne-modified 96-well plates. However, we found higher background for the ELLAs and ELISAs than for the dot blot assays (see SI). We also examined a commercial antibody with listed specificity to Sia, and lectins reported have linkage-specific preferences for Sia (Maakia amurensis I for α2,3Sia and Sambucus nigra I for α2,6Sia) but these exhibited poor specificity and high background.

Overall, we found that our sialylated and fucosylated synMUCs are indeed substrates for binding by natural glycan-targeting proteins. These data confirm the utility of our synMUCs as surrogates for native mucins, and as such, we expect these polymers to find broad utility in probing the diverse and fascinating biology of mucins. Further, considering the widespread use of glycan-binding lectins and antibodies in glycobiology research, we expect the synMUCs to be useful as standards or controls since the glycosylation can be controlled with exquisite precision.

### Conformational analysis of synMUC glycoforms

We utilized circular dichroism spectroscopy (CD) to examine the glycan-dependent secondary structures of our synMUC panel. This technique relies on the absorption of circularly polarized light by peptide bonds. The wavelengths at which the absorptions occur, and their intensities, reveal characteristics about the orientation of those bonds and the conformation of the peptide. Distinct CD signatures have been shown for the η→π* and π→π* amide bond transitions of disordered, β-sheet, α-helical, or polyproline II (PPII) helical conformations.^115,116^ SynMUC CD spectra were obtained in Milli-Q water and data were converted to molar ellipticity (Figure 5D–F). Homopolymers were utilized to hone in on the effects of glycosylation at Ser/Thr without convoluting effects of other amino acids.

Data in Figure 5D–F indicate that peptide conformation is strongly dependent upon glycan identity and amino acid linkage type. We observed a range of structures including α-helix, random coil, and extended or PPII. We note that the CD spectra of PPII helices (left-handed, 3 residues per turn), extended rod-like structures typical of polyelectrolytes, and disordered “random coil” conformations are quite similar. In fact, for many decades all of these structures were mistakenly assigned as random coils.^117,118^ Even now, despite their ubiquity in nature, PPII reference data are still absent from common deconvolution algorithms.^119–121^ However, careful comparison of the spectra of collagen, denatured collagen, intrinsically disordered proteins, and polyelectrolytes have shed light on the relationship between secondary structure and spectral absorbance wavelengths and intensities.^117,118,122–125^ Of particular focus is the η→π* region between ca. 205– 220 nm. Both polyanionic polyGlu and intact collagen have positive maxima, with the relative absorbance intensity increasing with degree of helicity or charge-repulsion-induced chain extension. By contrast, intrinsically disordered proteins have no positive maxima; absorbance is negative. The GalNAc amide contributes a positive absorbance between 190–205 nm in the π→π* region.^39,41^

Fig 5D shows an overlay of spectra obtained for various chemically-synthesized glycosylated polySer vs. polyThr backbones. Based on the η→π* transition intensity, we observed that chains of galactosylated-Thr are more extended, rigid, and PPII-like than those of galactosylated-Ser, likely due to steric contributions from the Thr methyls. Similarly, GalNAc results in more rigid, extended structures than Gal regardless of linkage to Ser vs. Thr. This is likely due to hydrogen bonding between glycan amides and the peptide backbone.^126^ Extension from GalNAc to the disaccharide Core 1 did not have a strong effect (see SI).

Sialylation had profound effects on glycopolypeptide secondary structure (Fig 5E). After addition of either α2,3Sia or α2,6Sia, to αGalNAc^S^ homopolymers, we observed distinct shifts in the spectra. We noted loss of the η→π* positive maximum at 220.8 nm and appearance of two new minima at ca. 212 and 223 nm, which is the classic signature of a canonical α-helix. Our data align with a prior report by Thornton and coworkers, who used CD and Raman techniques to characterize native MUC5B and observed helical and extended/disordered structures.^127^ To check if the drastic conformational shift is unique to our synthetic materials, we also collected CD spectra of bovine mucin before and after treatment with sialidase. Again, we observed an increased helical character with increased sialylation (see SI). Therefore, we conclude that sialylation induces mucin α-helicity. Uncovering the molecular drivers of this shift warrants detailed analyses in a future study. However, we hypothesize that the Sia carboxylic acid group could form a glycan-glycan hydrogen bond with a NAc group, and/or the steric bulk of the hydrated glycerol unit on Sia could disrupt the αGalNAc-to-peptide hydrogen bond which would free the peptide units for energetically favorable internal hydrogen bonds in the helix.

Fucosylation had less dramatic impacts on synMUC structure. All of the fucosylated βGal^T^ glycopolypeptides maintained the rigid, extended, PPII-like conformations observed for the monosaccharide-bearing precursors (Fig. 5C). We observed slight shifts in the wavelength of the η→π* positive maxima from 220.8 nm for βGal^T^ to 218.2, 216.1, and 215.6 nm for α1,2-, α1,3-, α1,4-Fuc modified βGal^T^, respectively. Following the same trend of increasing energy of absorbance wavelength, the magnitude of the absorption also increased with conjugation site. Together, these data suggest that peptide chain extension and rigidity is highest when Fuc is conjugated to C4 of the Gal ring, that fucosylation at C3, and even more so C2, results in slightly more flexible chains. Effects were greater for the Ser-based glycopolypeptide. For α1,2Fuc-βGal^S^, we noted loss of the η→π* maxima and shift to a spectrum analogous to that of truly disordered and flexible structures. Presumably, the relative hydrophobicity of the Fuc moiety and its proximity to the peptide backbone based on Gal linkage orientation, all affect the resulting structure. Again, the Thr methyl group appears to play a dominant role since Ser conjugates were more affected.

Collectively, the results of the structural analysis have implications for understanding evolution of mucin structure, interactions between mucins and their partners, as well as biophysical functions of the glycocalyx and mucus. Variations in glycoprotein secondary structures would result in differing presentations of glycans via their orientation in single or multivalent displays. Indeed, there are multiple reports of differences in the ability of antibodies and lectins to bind αGalNAc (Tn antigens) presented on Ser vs. Thr peptides.^77,128,129^ Considering that mucins are candidates for cancer vaccines and anti-infectives, understanding such effects is crucial. Further, the mechanics of the glycocalyx and mucus are intertwined with their biological roles and are affected by the rigidity and persistence length of component glycoproteins.^130–132^ Our data indicate that chain conformation and rigidity in mucin-type structures is glycan- and amino acid-dependent and is the likely result of an intricate balance of steric and hydrophobic effects, charge repulsion, and hydrogen bonding.

### Mucinase degradation

Proteolytic degradation of mucins occurs naturally in mucosal epithelial tissues that are colonized by microbes. Secreted mucinase enzymes play a variety of roles for commensal and pathogenic microbes alike, including in nutrition and infection. To shed light on the role of glycosylation in mucin degradation, we subjected our panel of synMUCs to two mucin-specific proteases of microbial origin. We chose secreted protease of C1 esterase (StcE), a zinc metalloprotease produced by *E. coli*, and *Serratia marcescens* Enhancin (SmE). StcE proteolysis occurs at Ser/Thr*-X-Ser/Thr sites, where cleavage occurs before the second Ser/Thr, X is any amino acid, and the (*) position must be glycosylated.^133^ SmE is reported to cleave *N*-terminally to a glycosylated Ser or Thr residue and between S/T-S/T sites.^73,134^ For both species, these mucinases contribute to cleavage of the protective mucus layers of the gut and, presumably, opportunistic infection.^135,136^ Further, these enzymes are under investigation for application as new research tools and therapeutics.^72,137,138^

We examined the susceptibility of various synMUCs to StcE and SmE degradation (Figure 6A). SynMUCs were incubated with the enzymes at an enzyme-to-substrate ratio of 1:10 for 48 hours at 37 °C, followed by heat denaturation of the mucinases. Control samples were exposed to the same conditions but without enzyme. Samples were subjected to sodium dodecyl sulfate polyacrylamide gel electrophoresis (SDS-PAGE) and stained with either silver stain or a glycoprotein-specific fluorescent stain. Representative gel data is shown in Figure 6B, C and complete gel data is in the SI. Gels were analyzed with ImageJ^139^ to quantify remaining intact polypeptide as compared to no-enzyme controls within the same gel.

**Figure 6.**
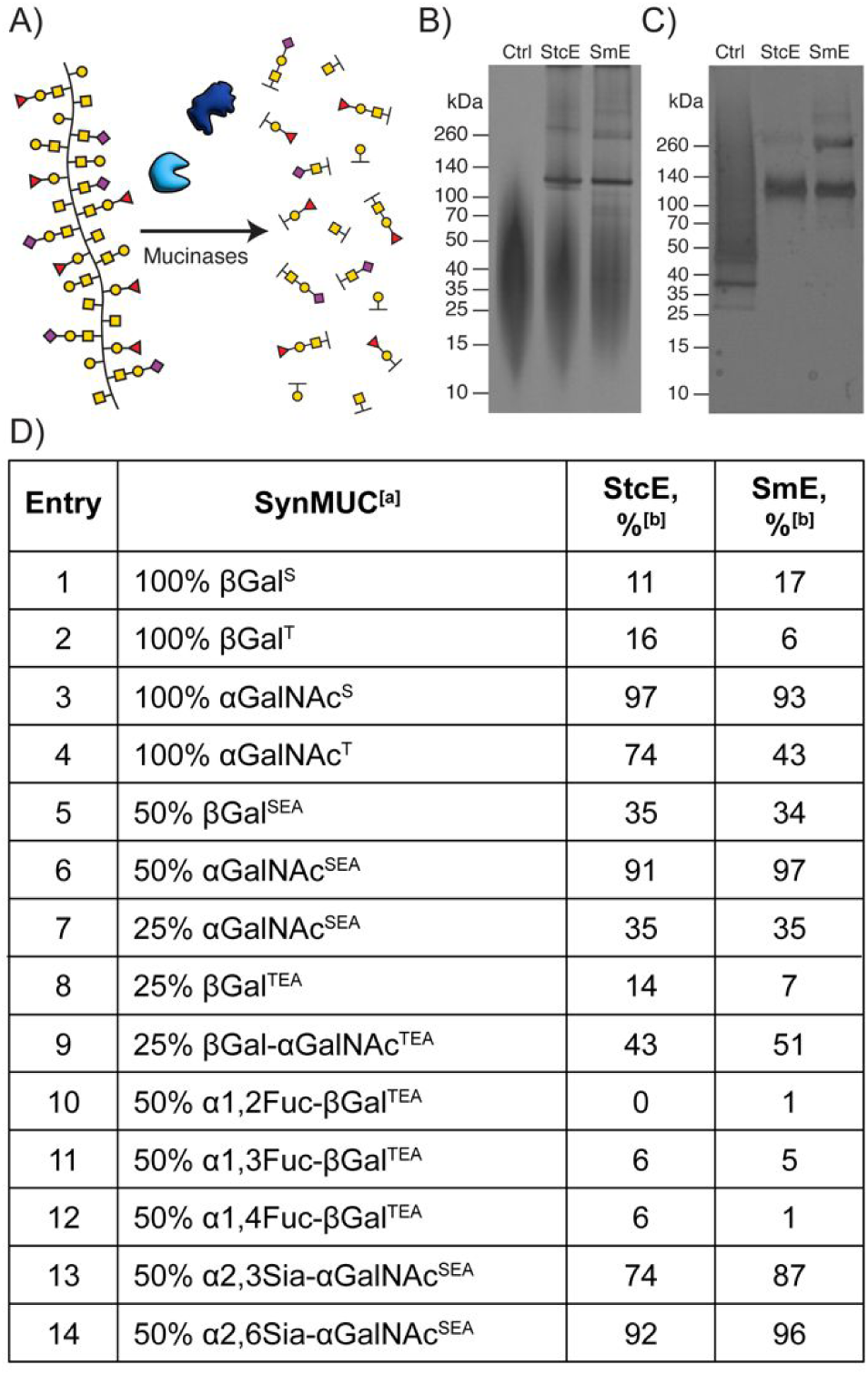
Degradation of synMUCs by mucinases StcE and SmE. A) Cartoon representation of the synMUC degradation experiment. B, C) Representative SDS-PAGE gels showing B) 50% βGal^SEA^, which was a poor mucinase substrate, vs. C) 100% αGalNAc^S^, which was an excellent mucinase substrate. In B, C) StcE and SmE mucinases appear at ca. 120kDa. D) Quantified percent degradation by image analysis of SDS-PAGE gels of mucinase treated synMUCs; [a] percentage denotes the glycosylation density, [b] denotes the percent degradation calculated by pixel density of the enzyme treated sample vs. the untreated control.

Both StcE and SmE preferred to cleave GalNAc-peptides as compared to Gal-peptides (Figure 6D Entries 1–9). We speculate the NAc group could have a stabilizing interaction within the active sites, or peptide secondary structure could more favorable. We also observed that proteolysis was more efficient for Ser vs. Thr conjugates of GalNAc (Figure 6D, Entries 3 vs. 4), and that SmE was more affected than StcE. We presume this is due to the higher steric constraints of the Thr methyl. Spacing of GalNAcylated residues to 50% did not affect proteolytic efficiency for either StcE or SmE, while spacing of Gal residues slightly increased proteolysis. Further decrease of glycan density to 25% reduced chain cleavage for both proteases over the time period examined, aligning with the fact that glycans are required for recognition by these enzymes (Figure 6D, Entries 3, 6, 7). Extension of the GalNAc residue to the disaccharide core 1 glycan did not appear to affect enzyme efficiency and the polypeptide was partially degraded during the time period examined (Figure 6D, Entry 9). Sialylation of GalNAc at C6 did not affect proteolysis by either StcE or SmE. Sialylation at C3 reduced proteolysis by 9% and 18% for SmE and StcE, respectively (Figure 6D, Entries 6 vs. 13 & 14) presumably due to altered access to the enzyme active site. By contrast, fucosylation of Gal at C2, C3, or C4 essentially eliminated the ability of both mucinases to cleave the peptide backbone.

Collectively, the mucinase data confirm our synMUCs can be recognized and degraded by natural enzymes and shed light on the proteolytic preferences of these two important enzymes. Prior mass spectrometry analyses eluded to glycan-dependent proteolysis by StcE and SmE; however, it was unclear if this was an artifact of detection methodology.^31,73,134^ SmE could accommodate a variety of *O*-linked glycans adjacent to the cleavage site, but might prefer smaller structures such as GalNAc or core 1 (Gal-GalNAc).^73,134^ StcE efficiently cleaved mucins bearing GalNAc or core 1, but sialylated-GalNAc mucins were reported as essentially resistant to proteolysis.^31^ By contrast, our data show that synMUCs bearing either α2,3 or α2,6-sialyated GalNAcs can be efficiently degraded. These data highlight the utility of chemically-defined mucin surrogates and their unique position to deconvolute structure-function relationships.

## Conclusions

Mucins are the primary protein component of mucus, saliva, tears, and the epithelial and endothelial glycocalyces. The heterogenous and dynamic nature of mucin glycosylation has been an obstacle to probing the biology of these tissues. We tackled this challenge by developing a chemoenzymatic route to prepare mucin-mimicking polypeptides. We used NCA polymerization to form glycopolypeptides bearing mono- and di-saccharides and then regio- and stereo-selectively diversified the glycans by enzymatic sialylation and fucosylation. After validation that these synthetic mucins are recognized by native glycan-binding proteins, we probed glycan-dependent effects on peptide backbone conformation and proteolytic susceptibility to mucinase enzymes. We found that glycosylation at Thr results in greater chain extension, and presumably rigidification, as compared to Ser residues. Similarly, GalNAc resulted in greater chain rigidification than Gal when linked at Ser or Thr. Fucosylation had only minor effects on secondary structure, while sialylation drove a conformation shift to alpha helices. We observed increased mucinase-catalyzed proteolysis for Ser- vs. Thr-based glycopolypeptides and for GalNAcylated vs. galactosylated and fucosylated structures. In summation, we present mucin mimics composed entirely of natural chemical groups and linkages, and that present multivalent displays of key bioactive fucose and sialic acid glycans. We expect these materials to serve as valuable tools for probing the diverse biology of mucins.

## Supporting Information

Full experimental details, additional data, and characterization of compounds can be found in the Supporting Information (SI).

## Acknowledgements

This work was funded by NIH NIGMS 1R35GM147262-01 to Jessica Kramer. We thank the lab of Prof. Michael Yu for the use of their CD spectrophotometer. We thank the lab of Prof. Russell Stewart for use of their aqueous SEC/MALS/RI system and Monika Sima and Dr. Rachel Detwiler for time and expertise on the related experiments. Finally, we thank Parastoo Azadi, Li Tan, and Christian Heiss at the Complex Carbohydrate Research Center for time and expertise with attempted analysis by mass spectrometry.

## TOC Graphic

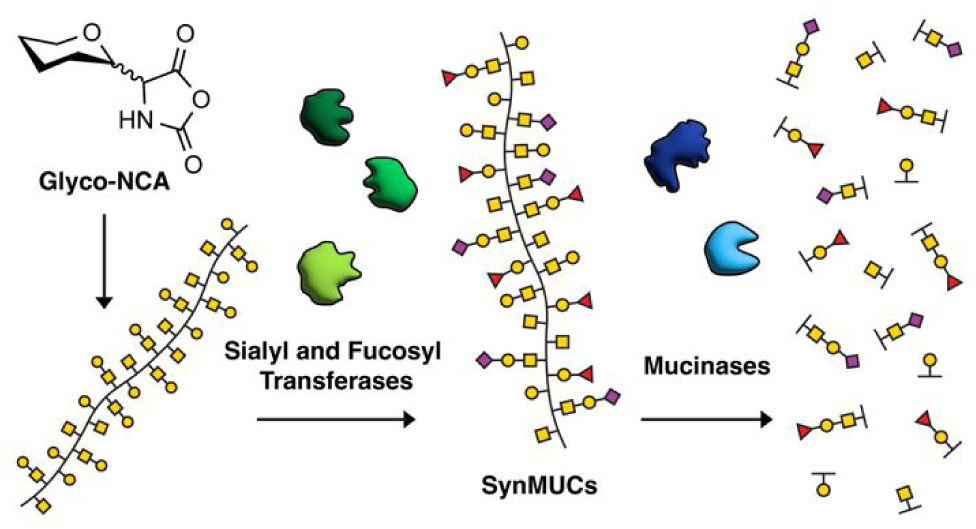

