## Supplemental Information for "Chemoenzymatic synthesis of sialylated and fucosylated mucin analogs reveals glycan-dependent effects on protein conformation and degradation" (from another version of this preprint: version 1, 2025-04-17)

### Table of Contents

**I. Materials and Methods**

General Instrumentation
Synthetic Mucin (SynMUC) Preparation
Enzyme Recombinant Syntheses
One-pot Multi-enzyme (OPME) SynMUC Glycosylations
ELLA/ELISA
Dot Blot
Circular Dichroism (CD)
X-ray Photoelectron Spectroscopy (XPS)
GC-MS and HPAEC-PAD
Mucinase Digestions and SDS-PAGE

**II. Supplementary Tables**

**S1** – Enzyme Recombinant Expressions
**S2** – Gibson Assembly Overlap Regions
**S3** – Restriction Enzymes
**S4** – Enzyme Purification Conditions
**S5** – GC temperature program for TMS glycosyl composition analysis
**S6** – HPAEC gradient used to separate sialic acid monosaccharides
**S7** – SynMUC Mucinase Digestion SDS-PAGE Parameters

**III. Supplementary Figures**

NMR
FTIR
GPC/MALS
CD
ELLA/ELISA
ELLA/ELISA Optimization
Dot Blot
GC-MS/HPAEC-PAD Analysis
XPS Spectra
Mucinase Digest Gels

**IV. Supplementary References**

### I. Materials and Methods

#### General Instrumentation

Anhydrous THF was purified by purging with dry nitrogen and then passing through columns of activated alumina. Ultra-pure water (18 MΩ-cm) was obtained by passing in-house deionized water through a Thermo Fisher Scientific MicroPure UV/UF purification tower. Infrared spectra were recorded on a Bruker Alpha ATR-FTIR Spectrophotometer to monitor all polymerizations. Protected SynMUCs were characterized via tandem gel permeation chromatography/light scattering (GPC/MALS/RI) with an Agilent 1260 Infinity liquid chromatography pump equipped with 18–angle Wyatt DAWN HELEOS-II light scattering (MALS) and Wyatt Optilab T-rEX refractive index (RI) detectors. SynMUCs were passed through $10^5$, $10^4$, and $10^3$ Å Phenominex Phenogel 5 μm columns in an eluent of 0.1 M LiBr in DMF at 60 °C at a concentration of 3 mg/mL. Enzymatically modified synMUCs and their deprotected precursors were further characterized via size exclusion chromatography coupled to a multi-angle light scattering and refractive index detector (SEC/MALS/RI). Analyses were performed on an Agilent PL aquagel-OH 30, 7.5 x 300 mm, 8 mm column, refractive index detector, and miniDawn TREOS detector (Wyatt Technology). SynMUCs were prepared at a concentration of 3 mg/mL and run in sterile PBS, pH 7.4 at a rate of 1 mL/min at 25 °C. ELLA/ELISA 96 well plate readouts were collected on a SpectraMax M2 microplate reader (Molecular Devices). Dot blots were imaged on an Azure 600 Biosystems imager with the IR-680 filter. Circular dichroism (CD) spectra were recorded in quartz cells with a path length of 0.1 cm on a JASCO J-1500 CD spectrophotometer. XPS was conducted in a Kratos Axis Ultra DLD X-ray/ultraviolet photoelectron spectrometer with an Al monochromator source at 1486.7 eV with anode voltage set to 15 eV and emission current set to 8 mA. All scans were performed under ultra-high vacuum with a pressure of greater than $3x10^{-9}$ Torr. A charge neutralizer was utilized and all C 1s spectra were charge corrected to 284.5 eV. GC-MS analysis was performed on an Agilent 7890A GC with a 5975C mass selective detector and 30 m x 0.25 mm inner diameter Supelco Equity-1 fused silica capillary column. HPAEC-PAD analysis was performed on a Dionex ICS3000 with a gradient pump, electrochemical detector, and autosampler. Separation was achieved with a 3 x 150 mm Dionex CarboPac PA20 column and amino trap guard column. SDS-PAGE gels were imaged with a UVP GelDoc-It imager.

#### Synthetic Mucin Preparation

Glyco-amino acid conjugates, *N*-carboxyanhydrides (NCAs), and polypeptides were prepared according to previously reported methodologies. *O*-(3,4,6-tri-*O*-acetyl-2-acetamido-2-deoxy-α-D-galactopyranosyl)-L-serine (αGalNAcSer) [1], L-alanine (Ala) [1], L-glutamic acid γ-*tert*-butyl ester (*t*BuGlu) [1], L-proline (Pro) [2], Nε-trifluoroacetyl-L-lysine (TFA-Lys)[3], *O*-((2,3,4,6,-tetra-*O*-acetyl)-β-D-galactopyranose))-L-threonine (βGalThr) [4] *O*-((2,3,4,6,-tetra-*O*-acetyl)-β-D-galactopyranose))-L-serine (βGalSer) [4,5], *O*-(2-acetamido-2-deoxy-3,4,6-tetra-*O*-acetyl-α-D-galactopyranosyl)-L-threonine, (αGalNAcThr) [4], and N-α-(butoxycarbonyl)-*O*-(2,3,4,6-tetra-*O*-acetyl-β-D-galactopyranosyl)-(1,3)-(2-acetamido-2-deoxy-4,6-*O*-acetyl-a-D-galactopyranosyl)-L-threonine (βGal-αGalNAcThr)[6] NCAs were prepared according to literature procedures. All polymerizations were performed in an $N_2$ filled glovebox. NCAs were suspended at 50 mg/mL in anhydrous THF and combined at various molar ratios to yield homopolymers or copolymers. Polymerizations were initiated at various monomer to initiator ratios via the rapid addition of $(PMe_3)_4Co$[7] or azido-Leu amido-amidate Ni[1] catalyst for polymers bearing a terminal azide (**Table 1**) at 30 mg/mL in dry THF. Polymerizations were allowed to proceed at RT in the glovebox or at 50 °C in a pressure vessel outside the glovebox for 3–72 hours (**Table 1**). Reaction progress was monitored via ATR-FTIR. Upon reaction completion, polymers were analyzed via GPC/MALS/RI. SynMUC protecting groups removed according to literature protocols[8] and all polymers were purified by dialysis or spin filtrations against 3kDa MWCO membranes in Milli-Q water, then lyophilized to obtain white powders in ca. 80–90% yields.

**Enzyme Recombinant Syntheses**

Each gene was cloned into the appropriate vector (**Table S1**) (Novagen, Addgene) via Gibson Assembly and designed in SnapGene software with overlap regions and restriction sites listed in **Tables S2** and **S3**. All enzymes were expressed with a poly-histidine tag for downstream purification. The plasmid (2 μg) was digested with the appropriate restriction enzymes (NEB, 40 U each) in NEB Cutsmart Buffer and the reaction proceeded at 37 °C for 60 minutes. The linear plasmid digest was purified via 0.7% agarose gel run at 100 V for 2.5 hours. The appropriate band size was excised and the plasmid recovered via gel purification (Zymo Gel DNA Recovery Kit). The Gibson Assembly of each plasmid was performed in 20 μL reactions with 0.02 pmol linearized plasmid and 0.06 pmol geneblock (IDT gBlocks) in Gibson Assembly Master Mix (NEB). Reactions were allowed to proceed at 50 °C for 60 minutes. Plasmids were immediately chemically transformed into TOP10 chemically competent E. coli cells (Thermo Fisher) via 30 second heat shock at 42 °C. Positive colonies were selected on carbenicillin agar plates grown for 17–20 hours at 37 °C and then grown up in Terrific Broth liquid culture with carbenicillin overnight to saturation. Positive clones were sequence verified before proceeding. Fresh broth was inoculated and grown with shaking at 250 RPM at 37 °C to an OD of 0.6–0.8 and then induced with IPTG at a concentration of 0.1 mM. Induced cultures were grown overnight with shaking at 225 RPM at room temperature. Bacteria were harvested via centrifugation at 4000xG for 30 minutes at 4 °C. The cell pellet was lysed with a buffer of 100 mM Tris-HCl pH 8.0 and 1% Triton-X at 20mL/liter culture. Lysozyme (VWR) was added at 100 μg/mL culture and DNase I (NEB) was added at 50 μg/liter culture. Cells were incubated at 37 °C for 50 minutes at 300 RPM and then centrifuged for 30 minutes at 12,000xG at 4 °C. Cell lysate was immediately purified via nickel or cobalt centrifugal affinity columns (Thermo Fisher) according to manufacturer protocol in a cold room or refrigerated centrifuge at 4 °C with buffer conditions described in **Table S4**. 5–8 elutions were taken of each enzyme and fractions with significant protein concentration were monitored via Nanodrop and combined. Purified enzymes were dialyzed at 4 °C against 50 mM Tris-HCl pH 7.5 and 10% glycerol with 4 buffer changes every 4–24 hours.[9,10] Enzymes were aliquoted and stored at -20 °C. Enzyme concentrations were quantified via BCA Assay (Thermo Fisher). Molecular weights were assessed via SDS-PAGE (Biorad, 4–12% Bis-Tris precast gels) ran in MES buffer (Biorad) at 150 V for 45 minutes and stained with GelCode Blue Safe Protein Stain (Thermo Fisher) or Pierce Silver Stain (Thermo Fisher).

**One-pot Multi-enzyme (OPME) SynMUC Glycosylations**

For all OPME glycosylation reactions, reagent equivalents were calculated per mole of Gal or GalNAc acceptor on the synMUC.

For all fucosylation reactions, 1 mg of synMUC was combined with 1.5 equiv. (per acceptor residue) L-fucose, 1.3 equiv. ATP, and 1.3 equiv. GTP in a buffer of 100 mM Tris-HCl, pH 7.5 with 20 mM $MgCl_2$ containing 0.1 mg/mL fucosyltransferase (Te2FT, HhFT1, or HpFT4), 0.067 mg/mL PmPpA, and 0.1 mg/mL BfFKP in a final volume of 500 μL. All concentrations listed are final concentrations. These conditions were scaled linearly with increased masses of synMUC. Fucosylation OPME reactions were modified from the work of Zhao et al.[11]

For all sialylation reactions, 1 mg of synMUC was combined with 1.5 equiv. (per acceptor residue) ManNAc, 7.5 equiv. sodium pyruvate, and 1.5 equiv. CTP in a buffer of 100 mM Tris-HCl, pH 7.5 with 20 mM $MgCl_2$ containing 0.2 mg/mL sialyltransferase (PmST1 or Pd26ST), 0.1 mg/mL NmCSS, and 0.2 mg/mL PmNanA in a final volume of 500 μL. All concentrations listed are final concentrations. These conditions were scaled linearly with increased masses of synMUC. Sialylations were allowed to proceed overnight at room temperature with gentle agitation at 100 RPM in an orbital shaker. Sialylation OPME reactions were modified from the work of Yu et al.[12]

For all completed SynMUC OPME reactions, polyhistidine-tagged enzymes were removed from the reaction mixture via incubation with NEBExpress Ni-NTA magnetic beads (NEB) according to manufacturer protocols. Finally, synMUCs were purified via dialysis against ultrapure water in 2 kDa tubing for 4 water changes every 4–24 hours or spin filtered with 3 kDa Amicon centrifugal filters, diluted 4 times in ultrapure water. SynMUCs were lyophilized to yield white fluffy powders.

**ELLA/ELISA**

Amine-coated 96 well plates (Bioworld) were incubated overnight at room temperature with a DBCO-NHS ester heterobifunctional crosslinker (Conju-Probe) (100 μL of 100 μg/mL in DMSO) with gentle rocking. Plates were washed 3 times with ultra-pure water to remove unbound crosslinker. DBCO coated plates were stored covered at 4 °C until use. Plates were coated with 50 μL of a 1 mg/mL solution of azide-terminal synMUC of interest in 1x PBS and left to conjugate at room temperature for 1–2 hours with gentle rocking. SynMUC conjugated plates were washed 3 times with PBST (1xPBS with 0.05% Tween-20) and used immediately for ELLA or ELISA.

For ELLA assays, 2 μg HRP-conjugated lectin of interest (EY Labs) was incubated in each well for 1 hour in PBS (100 μL final volume) at room temperature with gentle rocking. After 3 gentle washes with PBST, QuantaBlu fluorescent peroxidase substrate working solution (Thermo Fisher) was added according to manufacturer instructions and fluorescent readout was acquired 20 minutes post-addition.

For ELISA assays, surface-conjugated SynMUC was blocked with freshly prepared sterile filtered 3% BSA in PBS for 90 minutes at 37 °C. After 3 PBST washes, primary anti-sialic acid rabbit antibody (Antibodies Online) was diluted 1:1000 in PBS and 100 μL/well was incubated overnight (16 hours) at 4 °C. 3 PBST washes were performed before secondary HRP-conjugated goat anti-rabbit antibody was diluted 1:5000 in PBS and incubated 100 μL/well was incubated for 1 hour at room temperature with gentle rocking. After 3 final gentle washes, peroxidase fluorescent readout was performed as in the ELLA assays 20 minutes post-addition.

**Dot Blot Protocol**

0.2 μm nitrocellulose membranes (Biorad) were placed on filter paper and marked with a pencil to create a grid. SynMUC samples or control reagents were dissolved in TBS at concentrations of 30, 15, or 1 mg/mL for the fucose-binding lectin blot or 10, 5, or 1 mg/mL for the sialic acid-binding lectin blot and dotted in 1uL drops onto the membrane in triplicate. The human breast milk control was prepared by diluting raw milk 1 in 100, 1 in 300 and 1 in 600 in TBS. Samples were allowed to dry for 30 minutes before washing 3 times with 0.1% TBST for 5 minutes each wash with enough buffer to cover the membrane. The membrane was then blocked with 5% BSA in TBST for 1 hour at ambient temperature with gentle rocking. The membrane was washed 3 times with TBST. Biotinylated lectin (WGA to probe sialic acid or UEA-I to probe fucose, Vector Laboratories) was diluted 1:1000 in 3% BSA in TBST and incubated with the membrane for 16 hours at 4 °C. After washing 3 times with TBST, streptavidin labeled IR680 (Licor) was diluted 1:5000 in 3% BSA in TBST and incubated with the membrane for 1 hour at ambient temperature with gentle rocking. TBST was exchanged for TBS and the membrane was dried and imaged with an Azure 600 imager. Blot images were analyzed in ImageJ to quantify fluorescence intensity via ROI selection and quantification of integrated density.

**Circular Dichroism (CD)**

SynMUCs were dissolved in ultra-pure water at concentrations between 0.25–0.5 mg/mL. All spectra were recorded on a Jasco CD Spectrophotometer as an average of 3 scans between 190–250 nm wavelengths at 25 °C after blanking against ultra-pure water. CD ellipticity values were normalized to molar ellipticity with the formula $[\theta] = (\theta)/(10 * L * c)$, where θ is ellipticity (mdeg), L is cuvette pathlength (cm), c is concentration (M), and [θ] has the units $\mathrm{deg} * \mathrm{cm}^2/\mathrm{dmol}$. Spectra were plotted as $[\theta] * 10^{-3}\ \mathrm{deg} * \mathrm{cm}^2/\mathrm{dmol}$. Sample concentration was calculated using Beer's Law, $\mathrm{A} = \varepsilon \cdot \mathrm{c} \cdot \mathrm{l}$, by UV-Vis at 214 nm and extinction coefficient $\varepsilon$ = 2200 $cm^{-1}M^{-1}$ on a SpectraMax M2 spectrophotometer in a quartz cuvette with path length of 1 cm.

**X-ray Photoelectron Spectrometry (XPS)**

SynMUCs were prepared for XPS by aliquoting 100 μg of sample and lyophilizing to a fluffy white powder. All surfaces, tools, and gloves were sterilized with isopropanol before coming into contact with the sample or mounting surface. Carbon tape was cut to size with a clean razor blade and appended to the instrument loading bar to afford the fixation of indium foil. Indium foil was used to minimize contribution of unwanted carbon containing compounds from the adhesive carbon tape. SynMUC was pressed into the indium foil with a sterile spatula and briefly blown with nitrogen to remove unfixed powder to prevent sample contamination. Samples were placed inside the XPS load lock and pumped overnight to ultra-high vacuum conditions, about $10^{-7}$–$10^{-8}$ Torr. On the day of analysis, the loading bar was transferred to the sample analysis chamber vacuum between $10^{-8}$–$10^{-10}$ Torr. After charge neutralizer settings were adjusted, a low-resolution survey scan was collected with a pass energy of 160 eV and dwell time of 100 ms, followed by high resolution scans of O 1s, In 3d, N 1s, and C 1s regions with pass energy of 40 eV and dwell time of 300 ms. Five sweeps were collected for N 1s and 1 sweep was conducted for all other high-resolution scans. Data was analyzed in CasaXPS software. Low resolution scans were used to determine relative elemental composition and high resolution scans were deconvoluted into their characteristic peaks to identify relative percents of each elemental species.[13] FWHM values of all species were constrained to a range of 0.2 of each other and peak positions were constrained to known binding energies of a particular species.[13,14] Glycosylation efficiency was calculated by comparing the normalized ratio of C-N/C-O : C=O from the unmodified synMUC to the enzymatically modified synMUC as compared to the theoretical ratio of a 100% enzymatically glycosylated synMUC.

**GC-MS and HPAEC-PAD Analysis**

Glycan composition analysis was performed at the Complex Carbohydrate Research Center at University of Georgia. SynMUC glycopolypeptides and monosaccharide standards were derivatized to *O*-trimethylsilyl (TMS) methyl glycosides as described previously[15] with subsequent composition analysis performed via gas chromatography mass spectrometry (GC-MS). 300 μg of each lyophilized synMUC glycopolypeptide was resuspended at 1 mg/mL into 300 μL of ultra-pure water. 100 μL of each was then transferred to a scintillation vial containing 2 μg of inositol as an internal standard. Samples were lyophilized again before heating with 450 mL of methanolic 1 M HCl at 80 °C for 16 hours, followed by solvent removal under a stream of nitrogen. Depolymerized monosaccharides were then re-acetylated with methanol, pyridine, and acetic anhydride before drying again. Tri-Sil (Thermo Fisher) was used to derivatize each sample for 30 minutes at 80 °C. TMS methyl glycosides were dissolved in hexanes after drying with nitrogen and transferred to a new vial. Samples were dried again and resuspended with 100 μL of hexanes, and 1 μL was injected into the GC-MS. Analysis was performed with the temperature gradient shown in **Table S5** under a 10:1 split ratio mode. For sialic acid analysis, 100 μg aliquots of SynMUCs containing sialic acid were resuspended in 60 μL with a 1x solution of neuraminidase A digest buffer before the addition of 6 μL of α-2-3,6,8,9-neuraminidase A (New England Biolabs). Reactions incubated overnight at 37 °C, followed by the addition of 300 μL of ethanol. Samples were incubated at 4 °C for 3 hours before centrifugation at 10,000xG for 10 minutes. Supernatants were collected and precipitate pellets washed 2 times with 200 μL of 85% ethanol. Supernatant and washes of each sample were combined and dried under nitrogen. Each sample was resuspended in 100 μL of ultra-pure water and centrifuged a final time at 10,000xG for 10 minutes. Supernatants were then analyzed via HPAEC analysis compared to a standard calibration equation generated by linear interpolation of the respective area units of 3 serially diluted concentrations of Neu5Ac. The eluent gradient is described in **Table S6**, where 20 μL of each sample was injected every 35 minutes.

**Mucinase Protocols and SDS-PAGE Gels**

Mucinases (StcE[16] and SmE[17]) were kindly gifted from the lab of Prof. Stacy Malaker. For mucinase digestions, 200 μg synMUC polymer was incubated with StcE, SmE, or no mucinase in 1x PBS at an enzyme to substrate ratio of 1:10 in a final volume of 60 μL for 48 hours at 37 °C. After denaturing the mucinases for 5 minutes at 95 °C, digestions were stored at -20 °C until they were visualized via SDS-PAGE. 40–90 μg of digest was loaded per well with 4x XT sample buffer (Biorad) into a Criterion XT 4–12% Bis-Tris protein gel (Biorad). Gels were run at 175 V for 40 minutes in MES buffer (Biorad) and stained with either a Pierce Silver Stain Kit (Thermo Fisher) or a Pro-Q Emerald 300 Lipopolysaccharide Gel Stain Kit (Molecular Probes). SynMUC mass per well and stain conditions from SDS-PAGE are described in **Table S7**. SynMUC degradation was quantified via ImageJ.

### II. Supplementary Tables

**Table S1 - Enzyme Recombinant Expressions**

| Enzyme | Abbreviation | Plasmid Vector | GenBank Accession |
|---|---|---|---|
| α1,2 fucosyltransferase[9] | Te2FT | pET15b | NC_004113 |
| α1,3 fucosyltransferase[18] | HhFT1 | pMAL-c4x | AAP76669α |
| α1,4 fucosyltransferase[19] | HpFT4 | pMAL-c4x | AY450598 |
| L-fucokinase, GDP-fucose pyrophosphorylase[9] | BfFKP | pET15b | NC_003228 |
| Inorganic pyrophosphatase[9] | PmPpA | pET15b | NC_002663 |
| α2,3 sialyltransferase[20] | PmST1Δ24 | pET22b+ | NC_002663 |
| α2,6 sialyltransferase[10] | Pd2,6STΔ15 | pET15b | AAK02272 |
| Sialic acid aldolase[10] | PmNanA | pET15b | WP_000224714 |
| CMP-sialic acid synthetase[20] | NmCSS | pET22b+ | WP_002215295 |

**Table S2 – Gibson Assembly Overlap Regions**

| Plasmid | 5' Overlap Region | 3' Overlap Region |
|---|---|---|
| pET15b | TATACCATGGGCAGCAGCCATCATCATCATCATCACAGCAGCGGCCTGGTGCCGCGCGGCAGC | GGCTGCTAACAAAGCCCGAAAGGAAGCTGAGTT |
| pET22b+ | GCGGATAACAATTCCCCTCTAGAAATAATTTTGTTTAACTTTAAGAAGGAGATATA | CACCACCACCACCACCACTGAGATCCGGCTGCTAACAAAGC |
| pMAL-c4x | AGCTCGAACAACAACAACAATAACAATAACAACAACCTCGGGATCGAGGGAAGGATTTCA | GGCACTGGCCGTCGTTTTACAACGTCGTGACTGGGAAAACCCTGGCGTTACCCAACTTAA |

**Table S3 – Restriction Enzymes**

| Plasmid | 5' Restriction Enzyme | 3' Restriction Enzyme |
|---|---|---|
| pET15b | NdeI | BamHI |
| pET22b+ | NdeI | XhoI |
| pMAL-c4x | EcoRI | HindIII |

**Table S4 – Enzyme Purification Conditions**

| Enzyme | Affinity Column | Binding Buffer | Wash Buffer | Elution Buffer |
|---|---|---|---|---|
| Te2FT | $Ni^{2+}$ | 5mM imidazole, 0.5M NaCl, 50mM Tris-HCl pH 7.5 | 50mM imidazole, 0.5M NaCl, 50mM Tris-HCl pH 7.5 | 200mM imidazole, 0.5M NaCl, 50mM Tris-HCl pH 7.5 |
| HhFT1 | $Ni^{2+}$ | 5mM imidazole, 0.5M NaCl, 50mM Tris-HCl pH 7.5 | 20mM imidazole, 0.5M NaCl, 50mM Tris-HCl pH 7.5 | 200mM imidazole, 0.5M NaCl, 50mM Tris-HCl pH 7.5 |
| HpFT4 | $Co^{2+}$ | 10 mM imidazole, 0.3M NaCl, 50mM sodium phosphate; pH 7.4 | 10 mM imidazole, 0.3M NaCl, 50mM sodium phosphate; pH 7.4 | 150 mM imidazole, 0.3M NaCl, 50mM sodium phosphate; pH 7.4 |
| PmPpA | $Ni^{2+}$ | 5mM imidazole, 0.5M NaCl, 20mM Tris-HCl pH 7.5 | 20mM imidazole, 0.5M NaCl, 20mM Tris-HCl pH 7.5 | 200mM imidazole, 0.5M NaCl, 20mM Tris-HCl pH 7.5 |
| BfFKP | $Co^{2+}$ | 10 mM imidazole, 0.3M NaCl, 50mM sodium phosphate; pH 7.4 | 10 mM imidazole, 0.3M NaCl, 50mM sodium phosphate; pH 7.4 | 50 mM imidazole, 0.3M NaCl, 50mM sodium phosphate; pH 7.4 |
| PmST1Δ24 | $Ni^{2+}$ | 5mM imidazole, 0.5M NaCl, 20mM Tris-HCl pH 7.5 | 20mM imidazole, 0.5M NaCl, 20mM Tris-HCl pH 7.5 | 200mM imidazole, 0.5M NaCl, 20mM Tris-HCl pH 7.5 |
| Pd2,6STΔ15 | $Ni^{2+}$ | 5mM imidazole, 0.5M NaCl, 20mM Tris-HCl pH 7.5 | 20mM imidazole, 0.5M NaCl, 20mM Tris-HCl pH 7.5 | 200mM imidazole, 0.5M NaCl, 20mM Tris-HCl pH 7.5 |
| PmNanA | $Ni^{2+}$ | 5mM imidazole, 0.5M NaCl, 20mM Tris-HCl pH 7.5 | 20mM imidazole, 0.5M NaCl, 20mM Tris-HCl pH 7.5 | 200mM imidazole, 0.5M NaCl, 20mM Tris-HCl pH 7.5 |
| NmCSS | $Ni^{2+}$ | 5mM imidazole, 0.5M NaCl, 20mM Tris-HCl pH 7.5 | 20mM imidazole, 0.5M NaCl, 20mM Tris-HCl pH 7.5 | 200mM imidazole, 0.5M NaCl, 20mM Tris-HCl pH 7.5 |

**Table S5** – GC temperature program for TMS glycosyl composition analysis.

| | Rate (°C/min) | Value (°C) | Hold Time (min) |
|---|---|---|---|
| Initial | | 80 | 2 |
| Ramp1 | 20 | 140 | 2 |
| Ramp2 | 2 | 200 | 0 |
| Ramp3 | 30 | 250 | 5 |

**Table S6** – HPAEC gradient used to separate sialic acid monosaccharides.

| Time (min) | 100 mM NaOH | 1 M NaOAc in 100 mM NaOH |
|---|---|---|
| 0 | 97% | 3% |
| 7 | 97% | 3% |
| 15 | 50% | 50% |
| 17 | 30% | 70% |
| 20 | 30% | 70% |
| 22 | 97% | 3% |
| 35 | 97% | 3% |

**Table S7 – SynMUC Mucinase Digestion SDS-PAGE Parameters**

| **SynMUC** | **Sample Mass (μg)** | **Stain** |
|---|---|---|
| 100% $\beta Gal^{S}$ | 80 | Pro-Q Emerald |
| 100% $\alpha Gal^{T}$ | 80 | Pro-Q Emerald |
| 100% $\alpha GalNAc^{S}$ | 20 | Silver |
| 100% $\alpha GalNAc^{T}$ | 105 | Silver |
| 50% $\beta Gal^{SEA}$ | 20 | Silver |
| 50% $\alpha GalNAc^{SEA}$ | 90 | Silver |
| 50% $\alpha GalNAc^{SKA}$ | 80 | Pro-Q Emerald |
| 25% $\beta Gal^{TEA}$ | 80 | Pro-Q Emerald |
| 25% $\alpha GalNAc^{SEA}$ | 60 | Silver |
| 25% $\beta Gal\alpha GalNAc^{TEA}$ | 20 | Silver |
| 25% $\alpha GalNAc^{SEAP}$ | 20 | Silver |
| 50% $\alpha 1,2Fuc\beta Gal^{TEA}$ | 80 | Pro-Q Emerald |
| 50% $\alpha 1,3Fuc\beta Gal^{TEA}$ | 80 | Pro-Q Emerald |
| 50% $\alpha 1,4Fuc\beta Gal^{TEA}$ | 80 | Pro-Q Emerald |
| 50% $\alpha 2,3Sia\alpha GalNAc^{SEA}$ | 70 | Silver |
| 50% $\alpha 2,6Sia\alpha GalNAc^{SEA}$ | 60 | Silver |

### III. Supplementary Figures

#### NMR

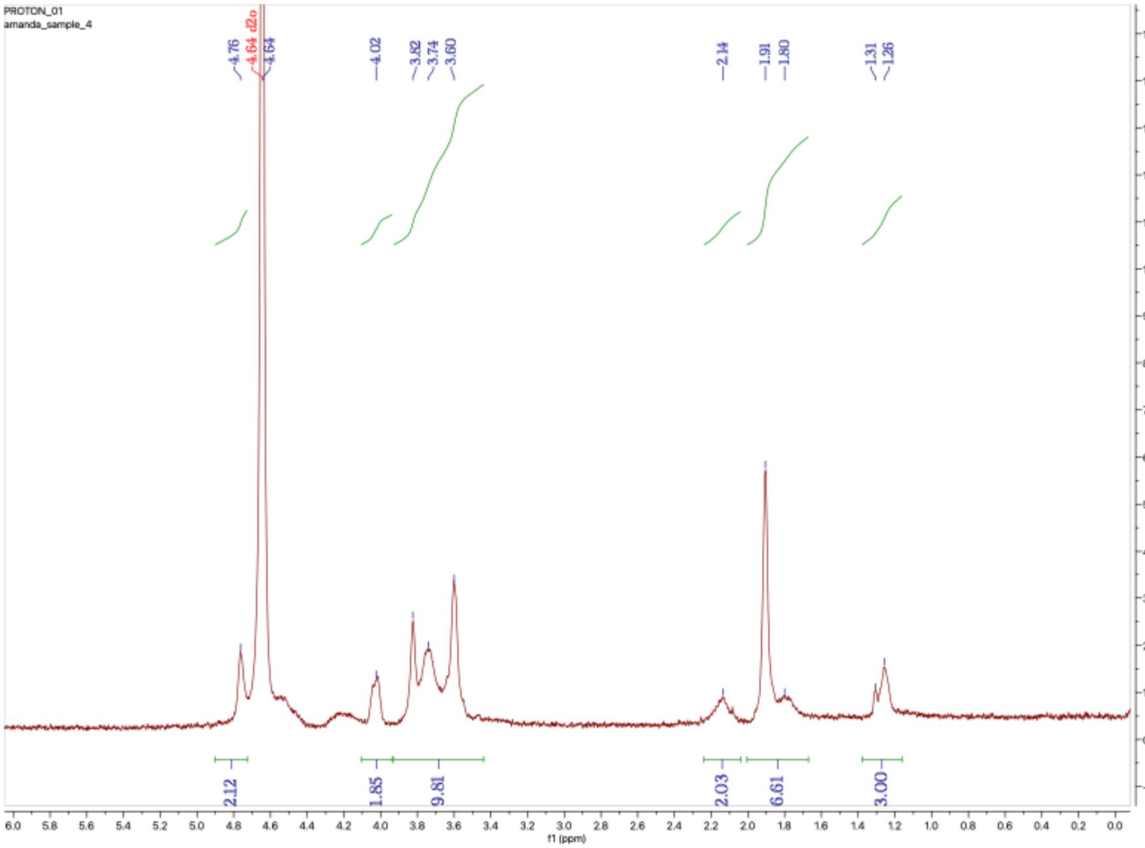


**Figure S1:** Representative $^1$H NMR ($D_2O$, 500 MHz). 50% αGalNAc$^{SEA}$, 98mer.

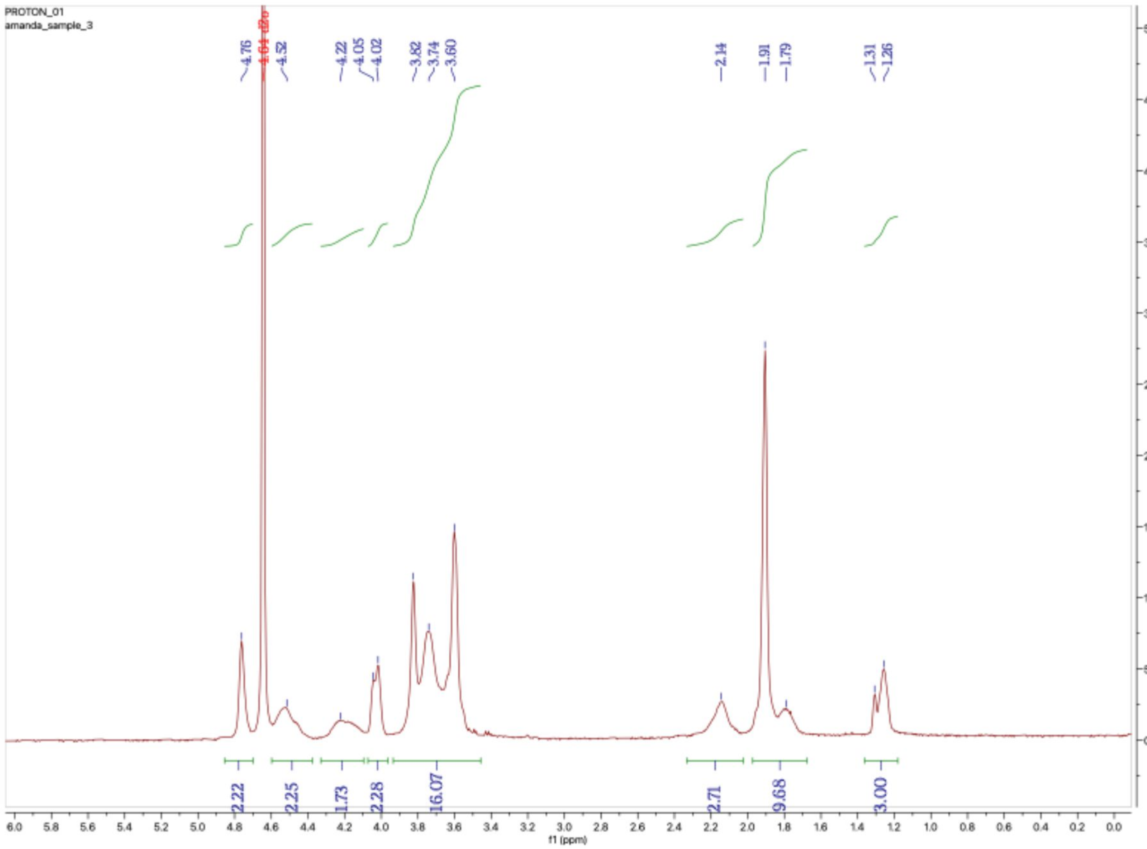


**Figure S2:** Representative $^{1}H$ NMR ($D_2O$, 500 MHz) of sialylated SynMUC. 50% α2,3Sia-αGalNAc$^{SEA}$, 98mer. Increased integration of proton signals between 1.65-1.95 ppm, ascribed to the sialic acid NAc methyl group, as compared to Figure S1 indicates enzymatic addition of sialic acid.

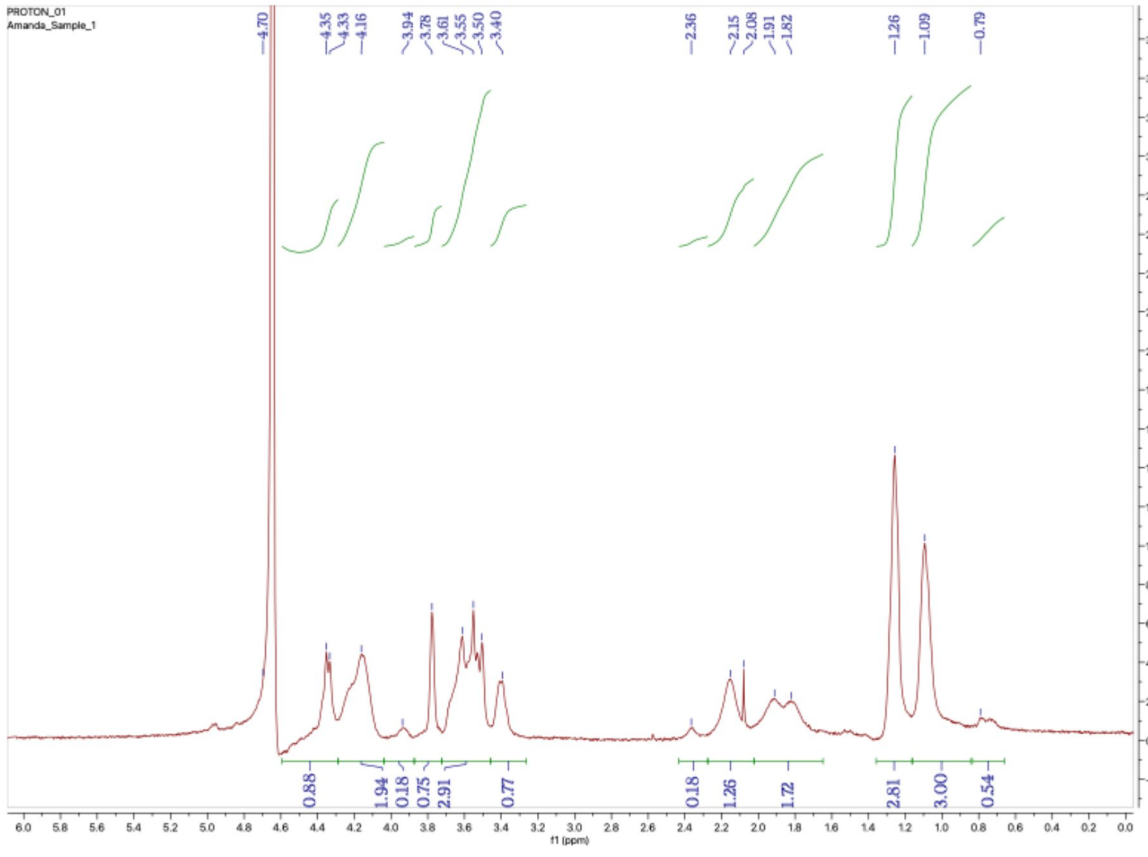


**Figure S3:** Representative $^1$H NMR ($D_2O$, 500 MHz). 50% βGal$^{TEA}$, 89mer.

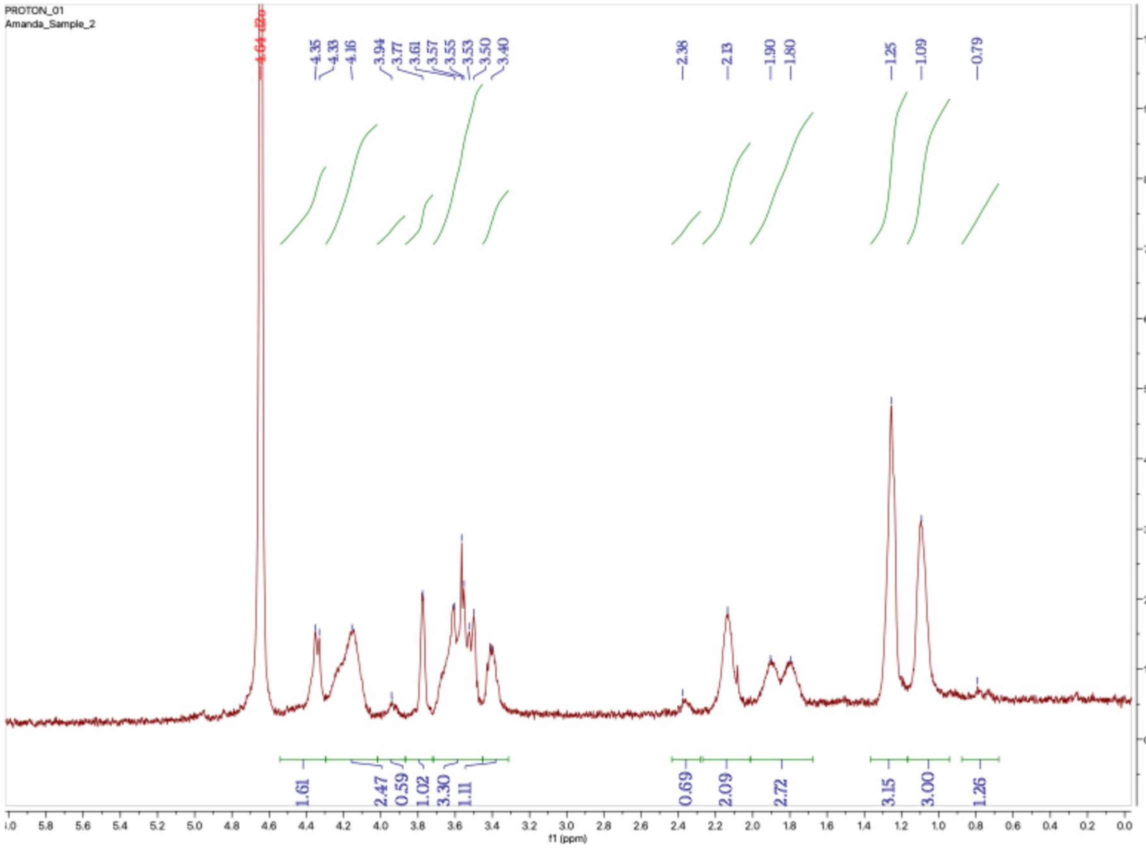


**Figure S4:** Representative $^1$H NMR ($D_2O$, 500 MHz) of fucosylated SynMUC. 50% α1,2Fuc-βGal$^{TEA}$, 89mer. Increased integration of proton signals between 1.15-1.35 ppm, ascribed to the fucose methyl group, as compared to Figure S3 indicates enzymatic addition of fucose.

**FTIR**

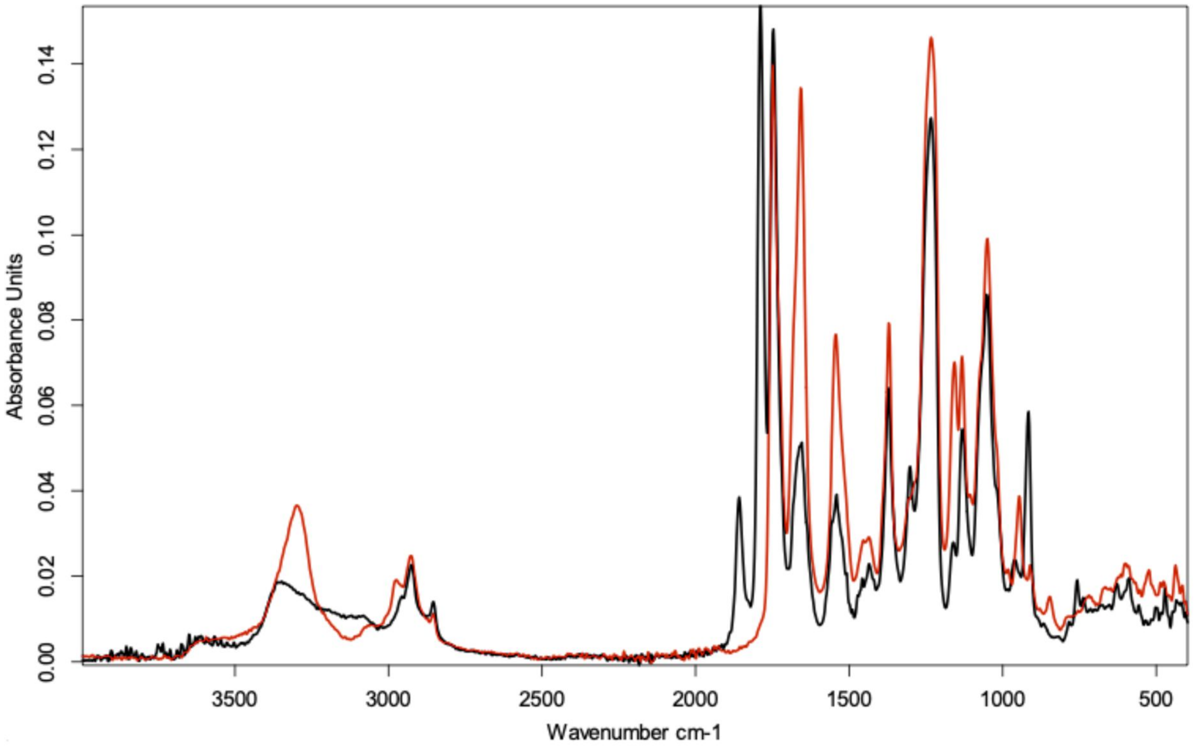


**Figure S5:** Representative FTIR spectra of αGalNAcSer NCA (black) overlaid with spectra of 50% αGalNAc$^{SEA}$ (red) at [M]:[I] = 30:1, indicating the complete consumption of NCA monomer and appearance of peptide bonds.

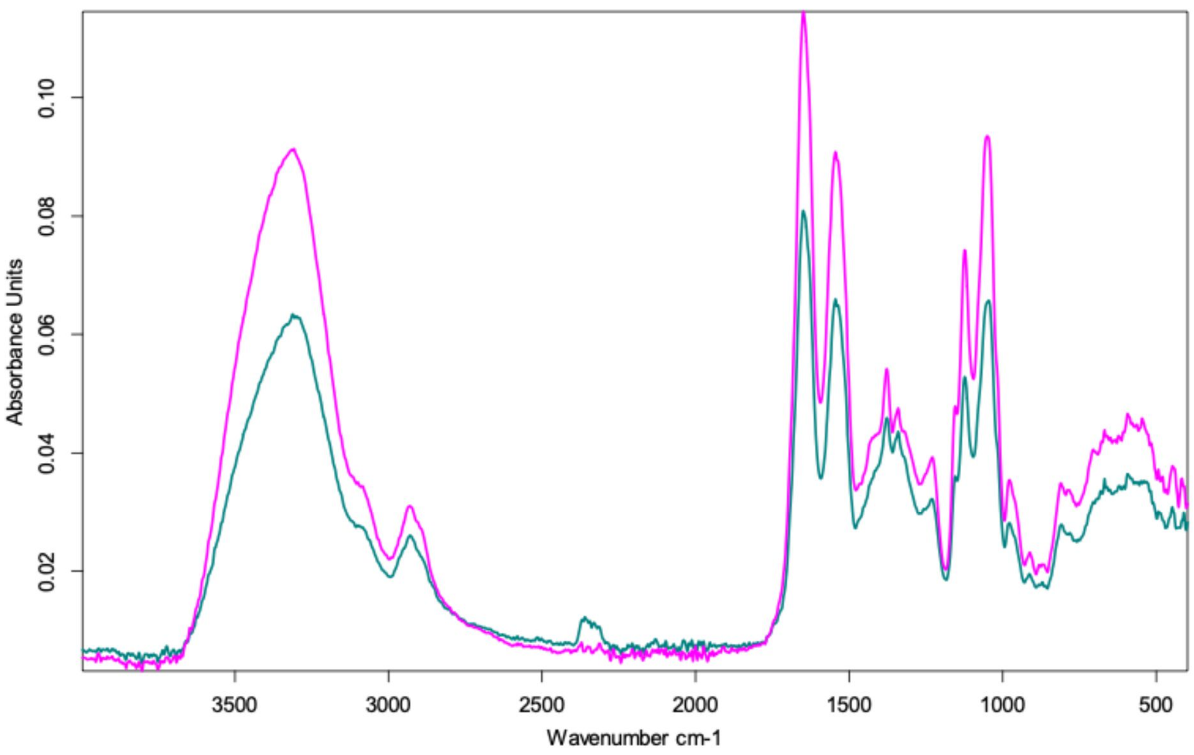


**Figure S6:** FTIR of the deprotected synMUC 100% αGalNAc$^{S}$, before (pink) and after sialylation α2,6Sia-αGalNAc$^{S}$ (teal).

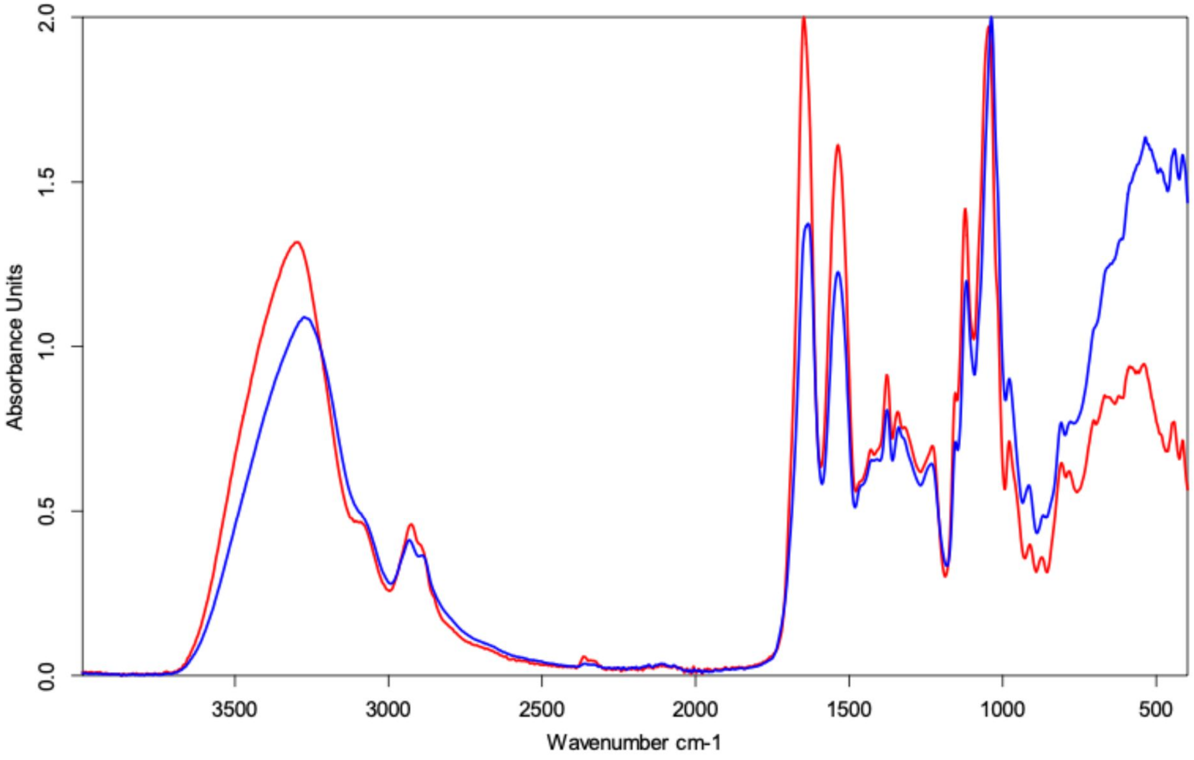


**Figure S7:** FTIR of the deprotected synMUC αGalNAc$^{S}$, before (red) and after sialylation α2,3Sia-αGalNAc$^{S}$ (blue).

### SEC/MALS/RI

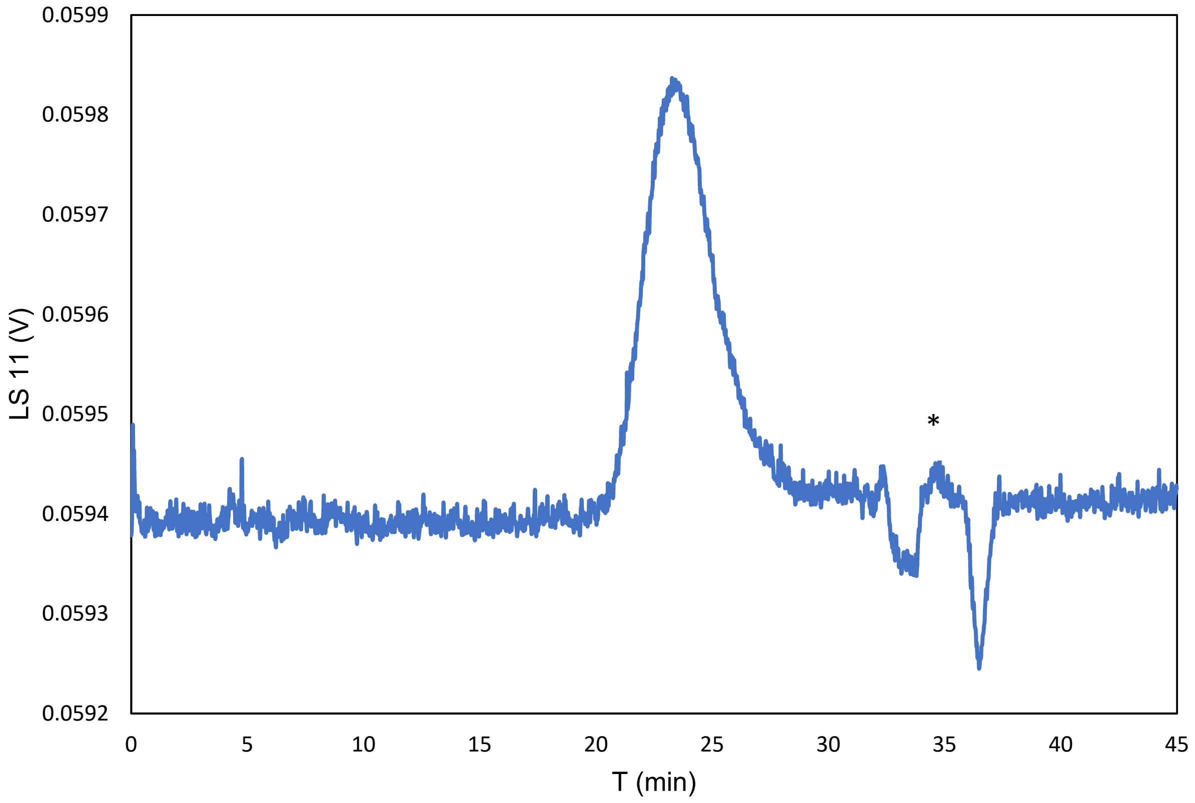


**Figure S8:** Representative SEC/MALS/RI (MALS trace shown) of 50% αGalNAc$^{SEA}$, [M]:[I] = 30:1, $Co(PMe_3)_4$ catalyst. *Denotes solvent peak.

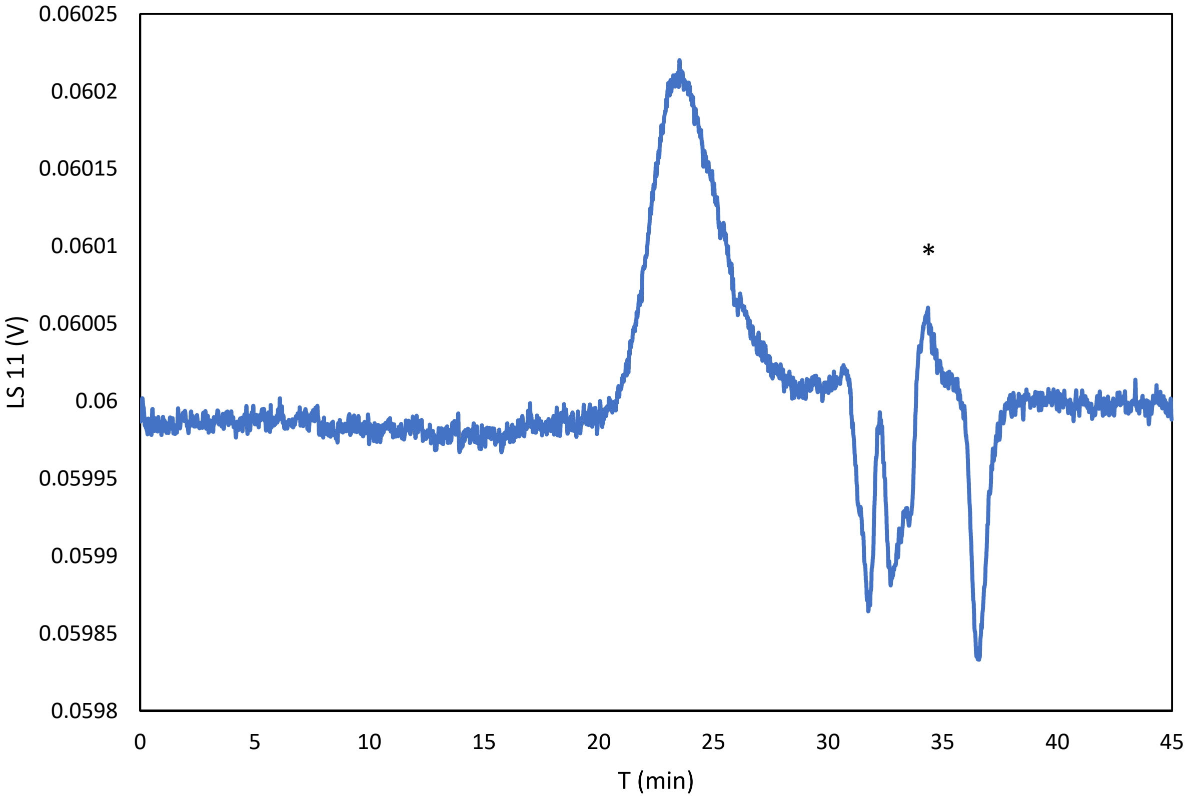


**Figure S9:** Representative SEC/MALS/RI (MALS trace shown) of 50% βGal$^{TEA}$, [M]:[I] = 30:1, Ni amido amidate catalyst. *Denotes solvent peak.

**CD**

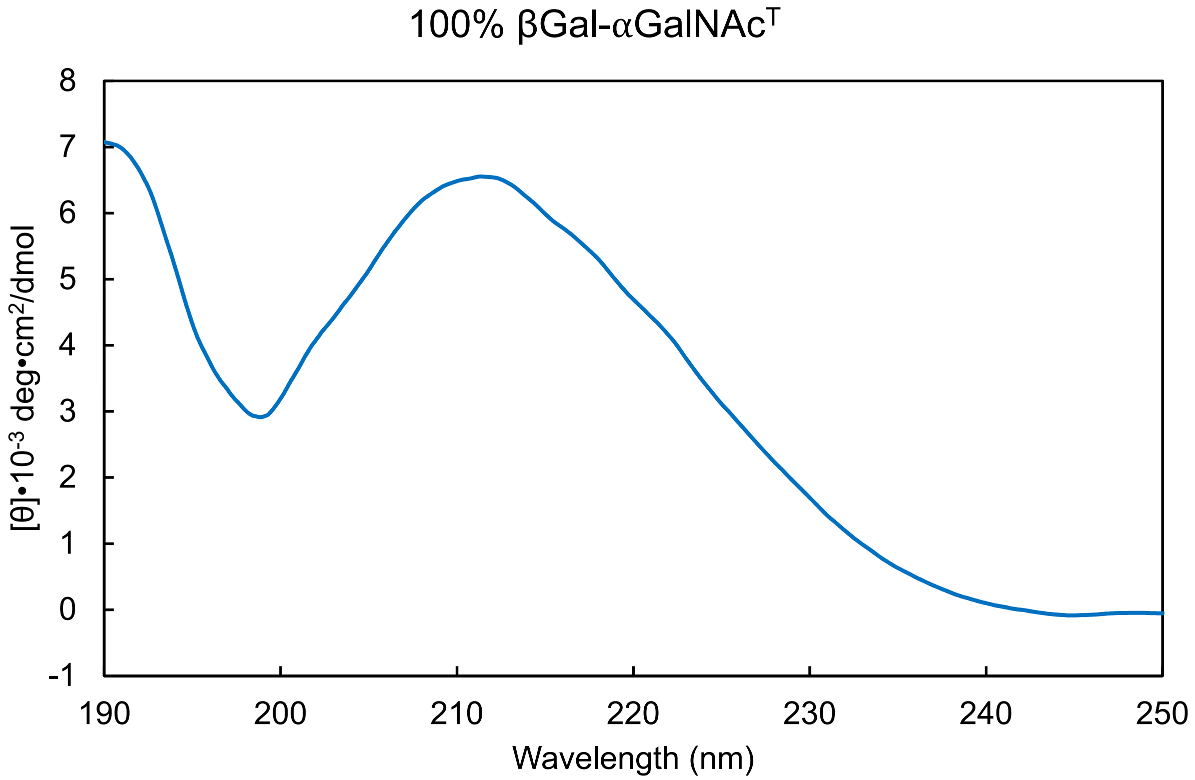


**Figure S10:** Circular dichroism spectra of a homopolymer of core 1 threonine.[21] Est. 75mer from [M]:[I].

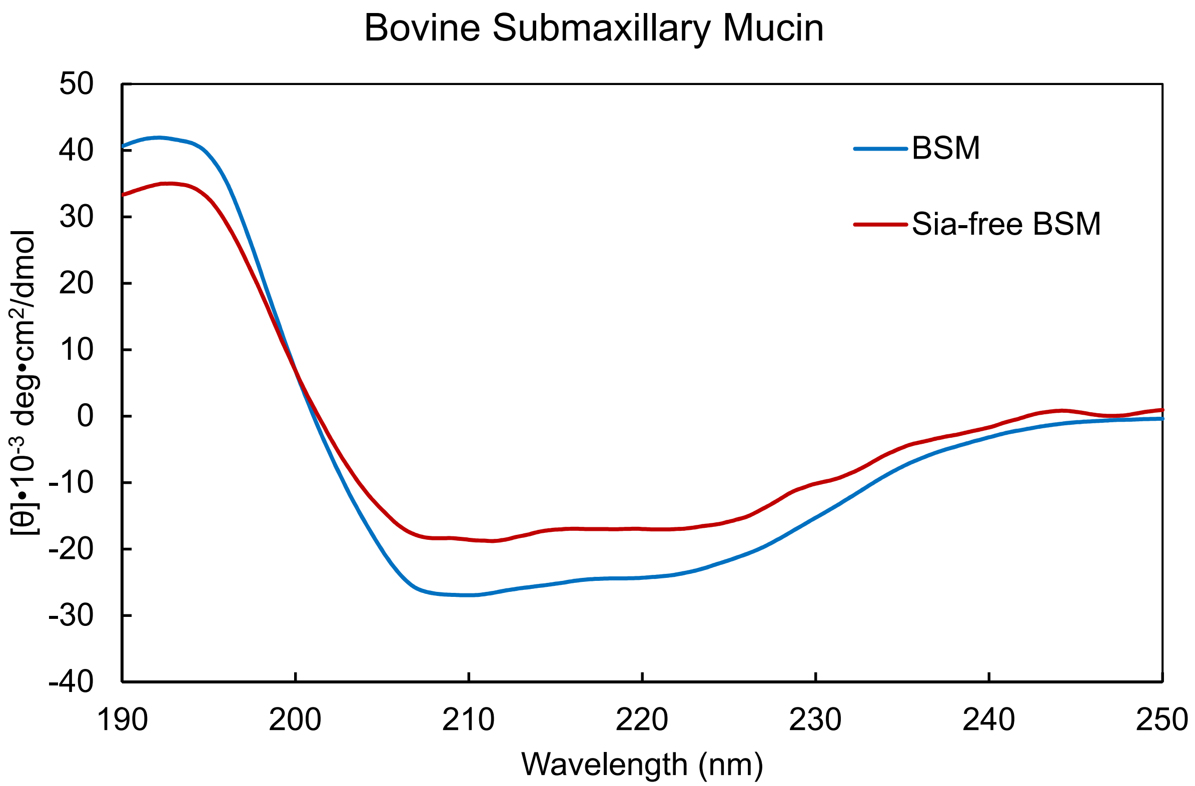


**Figure S11:** Circular dichroism spectra of bovine submaxillary mucin (BSM) with and without sialic acid. BSM with sialic acid has stronger alpha helical character than BSM treated with neuraminidase.

**ELLA/ELISA**

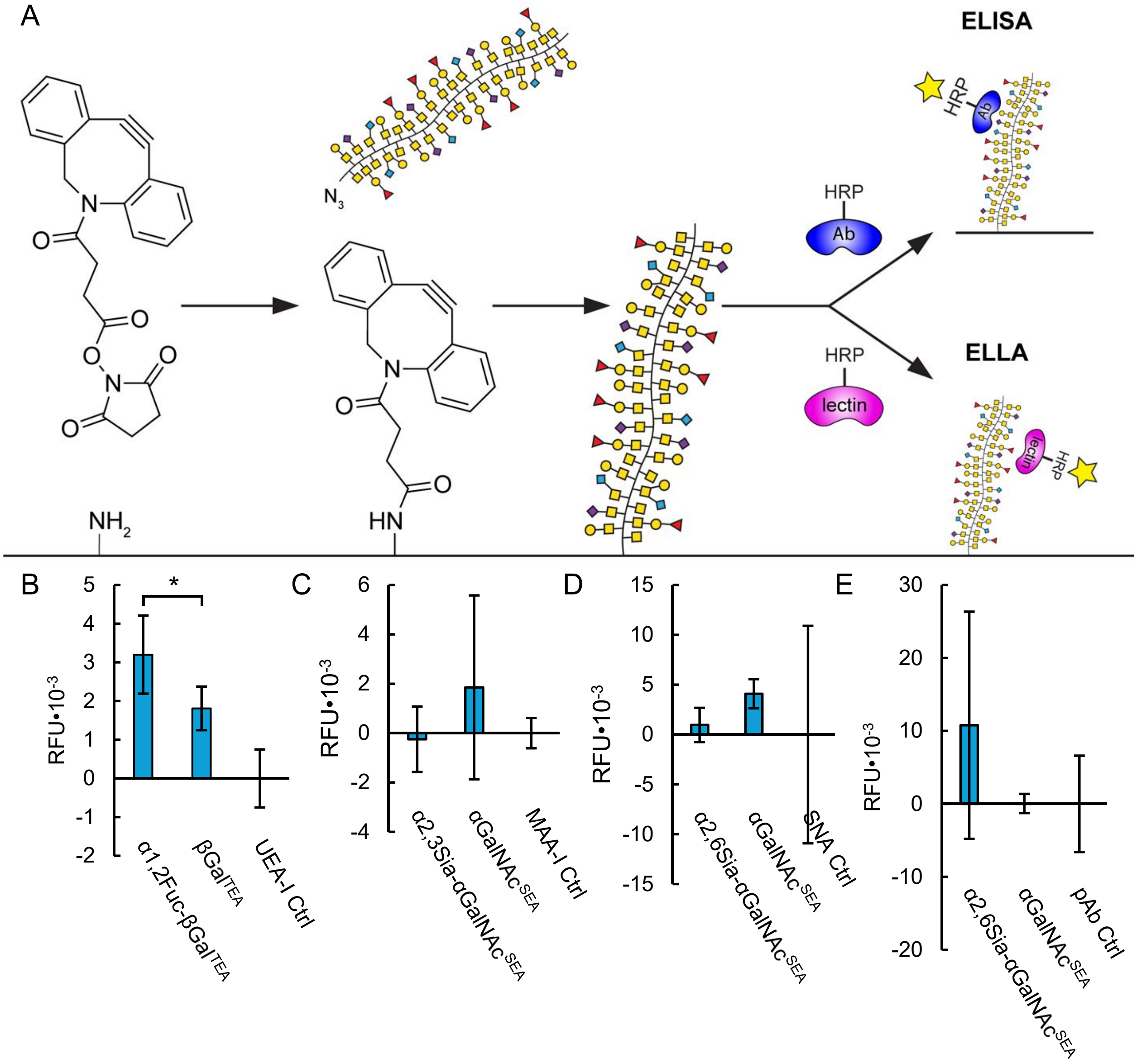


**Figure S12**: ELLA and ELISA based recognition of synMUCs with the terminal glycans fucose or sialic acid. A) Experimental workflow of ELLA and ELISA experiments. An amine-coated multi-well plate was incubated with a heterobifunctional crosslinker NHS ester–DBCO. An azide-terminated SynMUC was then introduced to covalently click the SynMUCs to the plate. HRP-labelled lectins or antibodies were then incubated with the samples and a fluorescent readout was collected. B) ELLA confirms presence of fucose on a SynMUC. SynMUC with and without fucose were tested under these assay conditions to show a significant difference in fluorescence before and after fucosylation. MAA-I ELLA (C), SNA ELLA (D), and sialic acid ELISA (E) all show inconsistent binding and high background. *$p<0.05$.

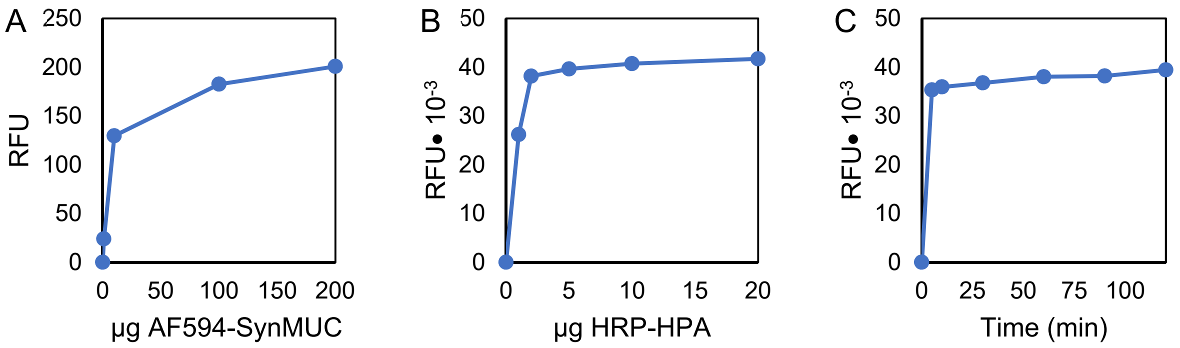


**Figure S13:** ELLA/ELISA optimization. A) Increasing masses of azide-terminal synMUC labelled with the fluorophore AF594 were tested on the DBCO conjugated 96 well plate to determine optimal mass of synMUC to use per well. 50 μg of synMUC was chosen for subsequent experiments. SynMUC used was azide-terminal 15% αGalNAc$^{SEA}$ , 200mer (60% E, 25% A). B) Increasing masses of lectin were incubated with synMUC to determine saturation point. 2 μg of lectin was chosen for subsequent experiments. SynMUC used was 50% αGalNAc$^{SEA}$, 185mer, and lectin was HRP conjugated Helix Pomatia Agglutinin (HRP-HPA). C) Finally, lectin incubation time was tested to determine signal saturation. 60 minutes of incubation time was chosen for subsequent experiments. SynMUC used was 100% αGalNAc$^{S}$, 25mer and lectin was HRP-HPA.

### Dot Blot

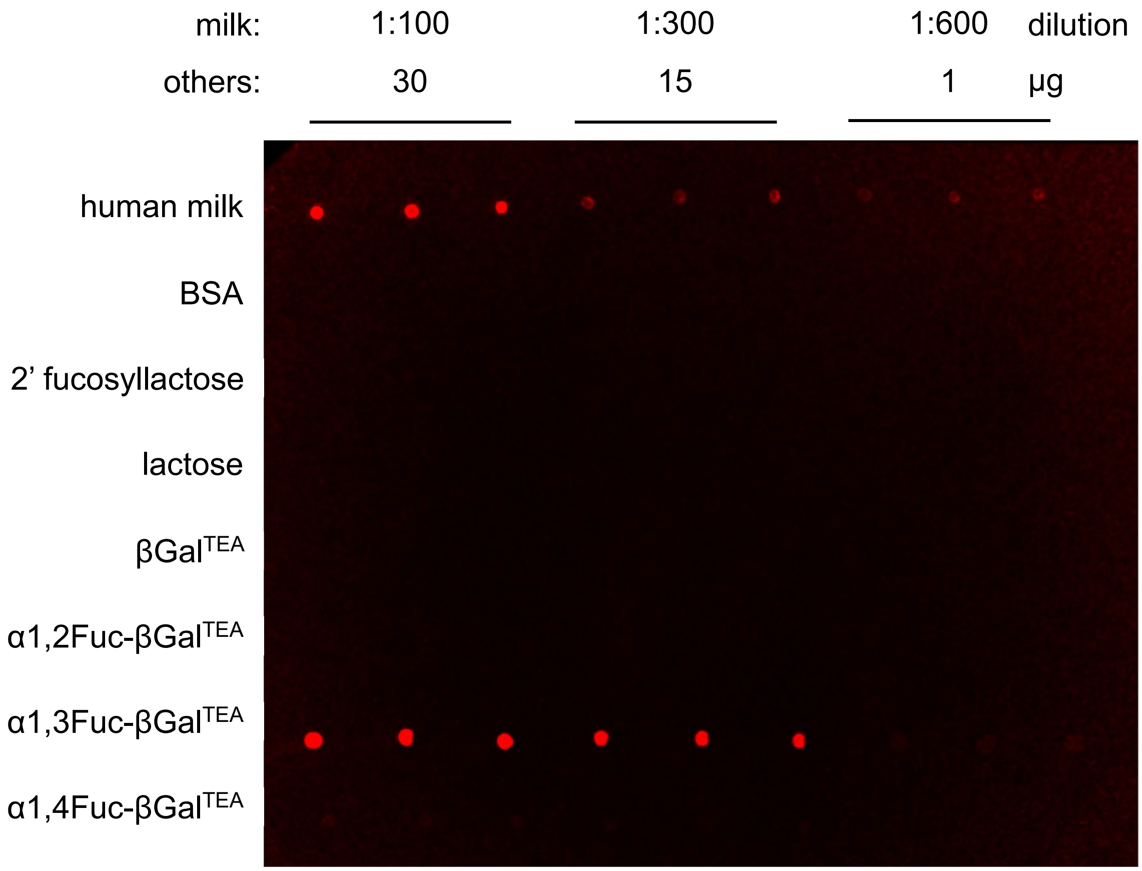


**Figure S14:** UEA-I dot blot of fucosylated SynMUCs. All samples dotted at 30, 15, and 1 µg and human milk dotted at a dilution of 1 in 100, 1 in 300, and 1 in 600 in TBS. 2' fucosyllactose was purchased from Sigma.

### GC-MS/HPAEC-PAD

| SynMUC Name | Mass Gal (mg) | Mass GalNAc (mg) | Mass Fuc or Sia (mg) | Moles Gal | Moles GalNAc | Moles Fuc or Sia | % Glyco Efficiency |
|---|---|---|---|---|---|---|---|
| 100% α1,2Fuc -βGal$^{S}$ | 28.27 | 0 | 1.37 | 1.57E-7 | 0 | 8.35E-9 | 5.32 |
| 100% α1,2Fuc -βGal$^{T}$ | 12.83 | 0 | 1.33 | 7.12E-8 | 0 | 8.10E-9 | 11. 38 |
| 100% α2,6Sia -αGalNAc$^{S}$ | 0 | 114.2 | 0.02 | 0 | 5.16E-7 | 6.47E-11 | 0.01 |
| 100% α2,6Sia -αGalNAc$^{T}$ | 0 | 105 | 0.04 | 0 | 4.75E-7 | 1.29E-10 | 0.03 |
| 25% α1,2Fuc -βGal$^{TEA}$ | 61.32 | 0 | 0.57 | 3.40E-7 | 0 | 3.47E-9 | 1.02 |
| 50% α1,2Fuc -βGal$^{TEA}$ | 34.76 | 0 | 0.11 | 1.93E-7 | 0 | 6.70E-10 | 0.35 |
| 50% α1,3Fuc -βGal$^{TEA}$ | 38.77 | 0 | 0.02 | 2.15E-7 | 0 | 1.22E-10 | 0.06 |
| 50% α1,4Fuc -βGal$^{TEA}$ | 46.31 | 0 | 0.03 | 2.57E-7 | 0 | 1.83E-10 | 0.07 |
| 25% α2,6Sia - αGalNAc$^{SEA}$ | 0 | 9.2 | 0.4 | 0 | 4.16E-8 | 1.29E-9 | 3.11 |

| | | | | | | | |
|---|---|---|---|---|---|---|---|
| 50% α2,3Sia -αGalNAc$^{SEA}$ | 0 | 30.8 | 0.01 | 0 | 1.39E-7 | 3.23E-11 | 0.02 |
| 50% α2,6Sia -αGalNAc$^{SEA}$ | 0 | 10.5 | 0 | 0 | 4.75E-8 | 0 | 0.00 |
| 25% α1,2Fuc -βGal$^{TEAP}$ | 53.29 | 0 | 0.59 | 2.96E-7 | 0 | 3.59E-9 | 1.22 |
| 25% α2,6Sia -αGalNAc$^{SEAP}$ | 0 | 65.3 | 0.95 | 0 | 2.95E-7 | 3.07E-9 | 0.03 |
| 50% α1,2Fuc -βGal$^{TKA}$ | 58.15 | 0 | 0.15 | 3.23E-7 | 0 | 9.14E-10 | 0.28 |
| 50% α2,6Sia -αGalNAc$^{SKA}$ | 0 | 74.4 | 0.03 | 0 | 3.36E-7 | 9.70E-11 | 0.03 |
| 25% α1,2Fuc -βGal -αGalNAc$^{TEA}$ | 24.41 | 0.763 | 0.74 | 1.35E-7 | 3.45E-9 | 4.51E-9 | 3.33 |
| 25% α2,6Sia -βGal -αGalNAc$^{TEA}$ | 18.9 | 37.9 | 2.38 | 1.05E-7 | 1.71E-7 | 7.70E-9 | 4.49 |
| 25% α1,3Fuc -βGal -αGalNAc$^{TEA}$ | 11.87 | 23.7 | 0.64 | 6.59E-8 | 1.07E-7 | 3.90E-9 | 5.92 |
| 25% α1,4Fuc -βGal -αGalNAc$^{TEA}$ | 4.58 | 8.2 | 0.21 | 2.54E-8 | 3.71E-8 | 1.28E-9 | 5.03 |

**XPS Spectra**

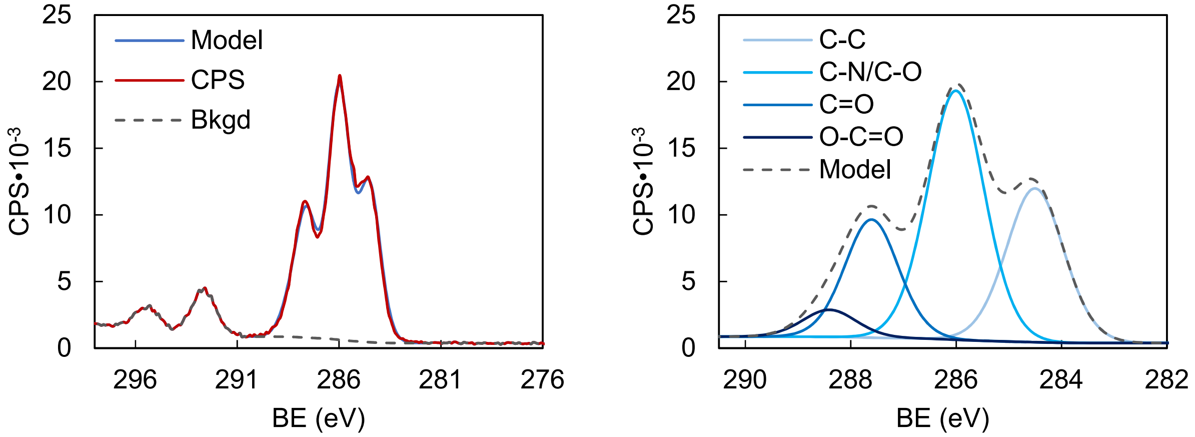


*50% αGalNAc[SEA]*

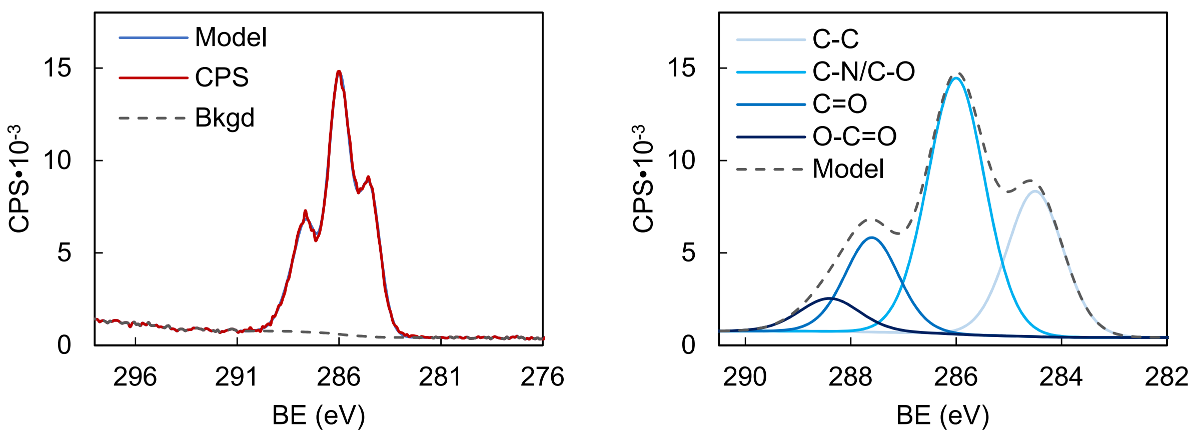


*50% α2,3Sia-αGalNAc[SEA]*

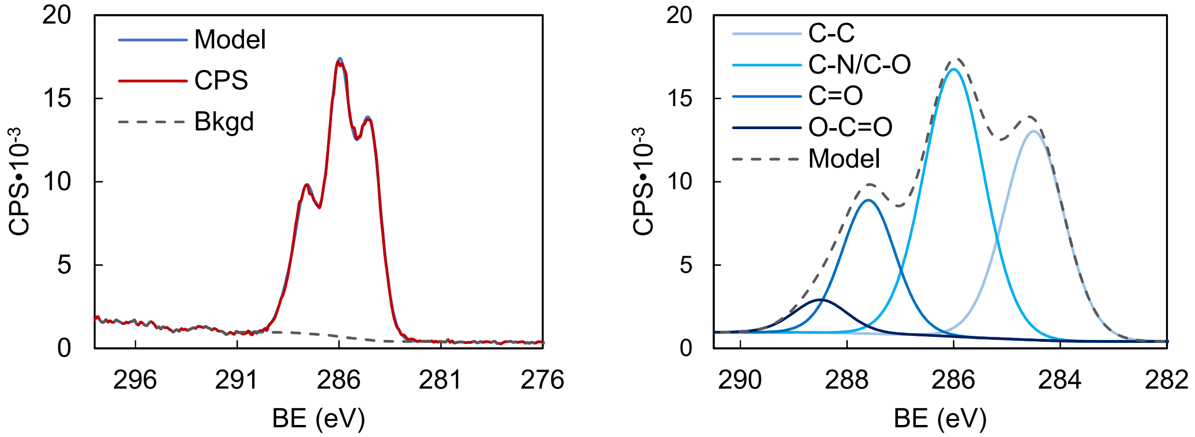


*50% αGalNAc[SEA]*

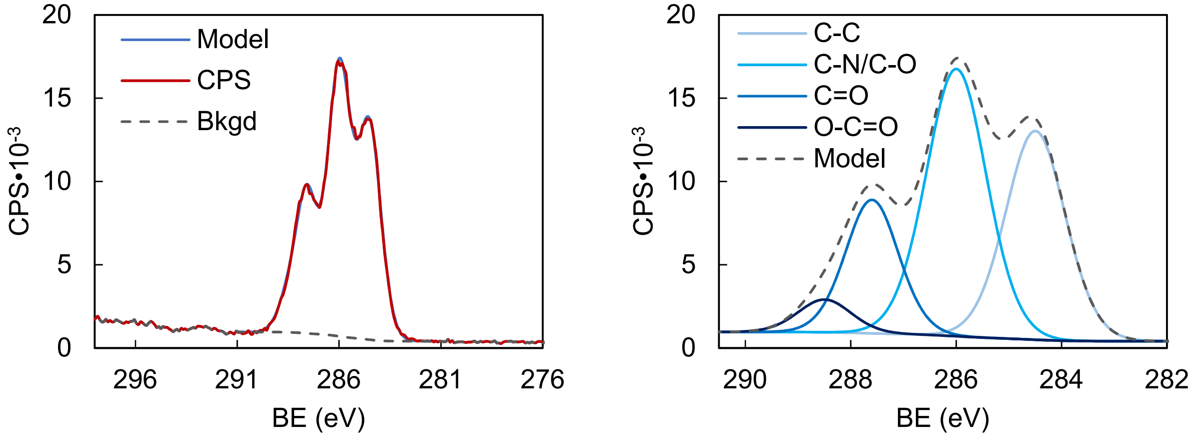


*50% α2,6Sia-αGalNAc*$^{SEA}$

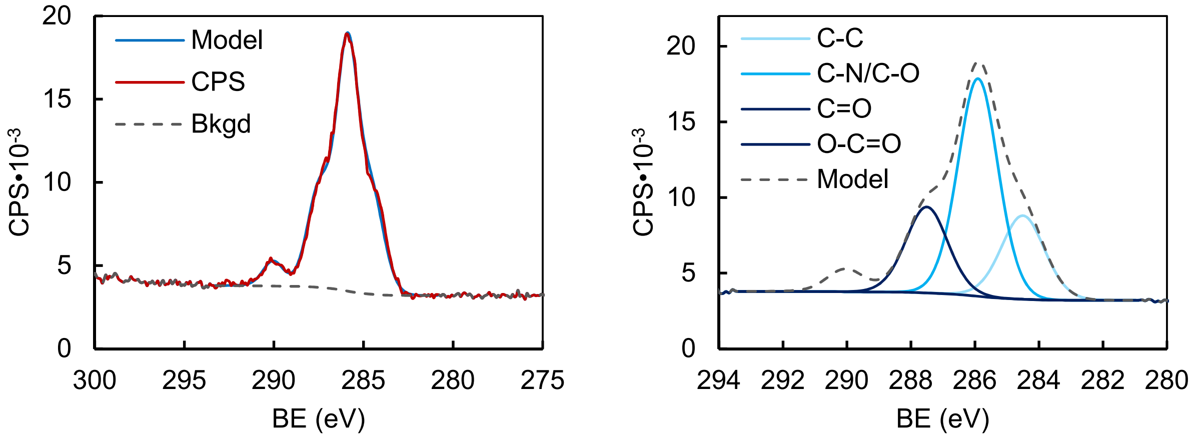


*50% αGalNAc*$^{SKA}$

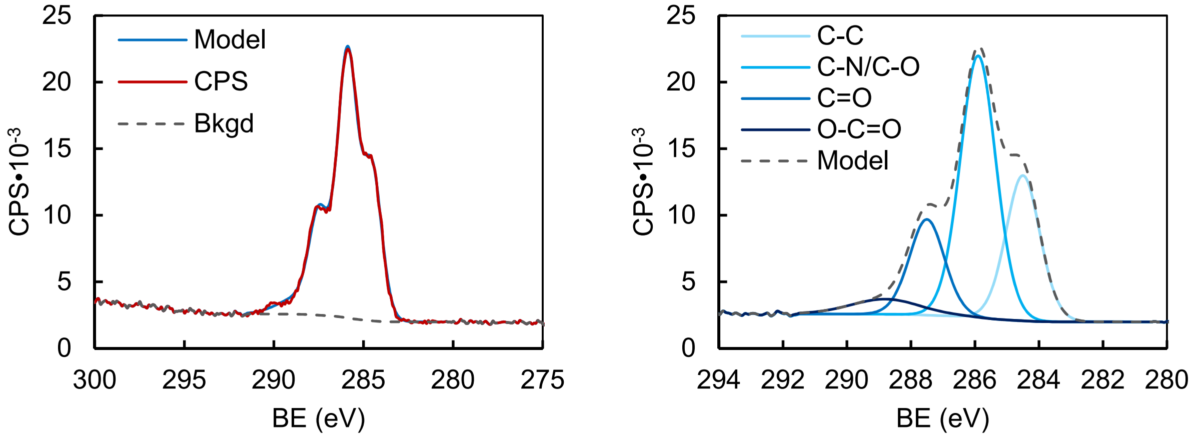


*50% α2,6Sia-αGalNAc*$^{SKA}$

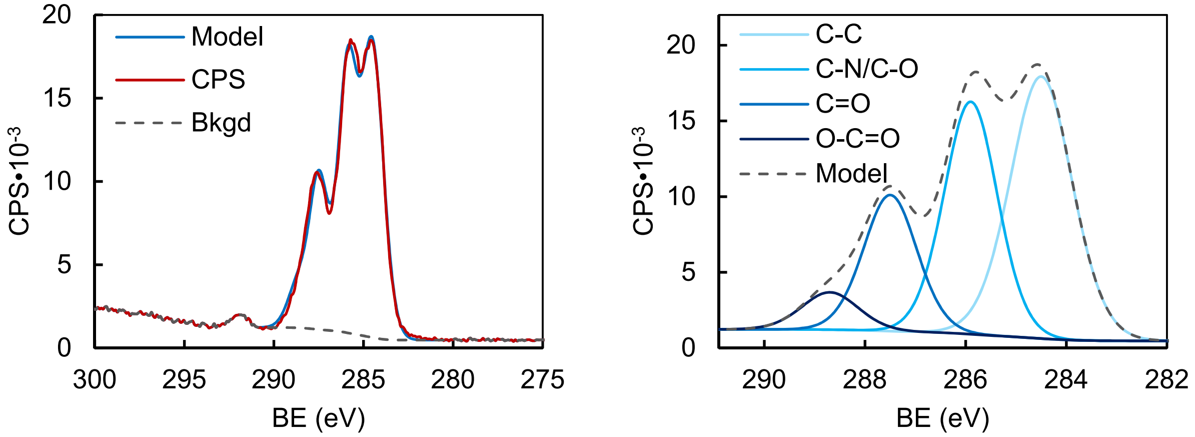


*25% αGalNAc$^{SEAP}$*

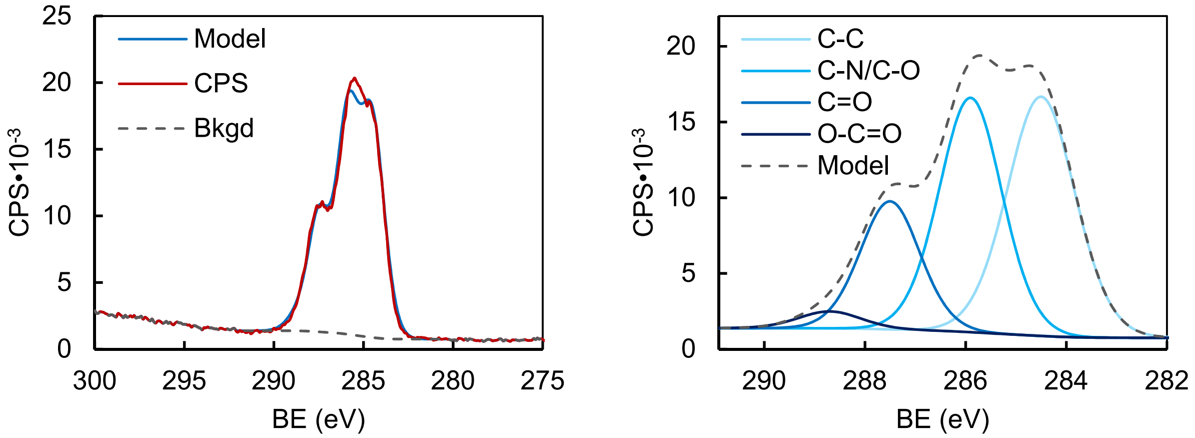


*25% α2,6Siaα-GalNAc$^{SEAP}$*

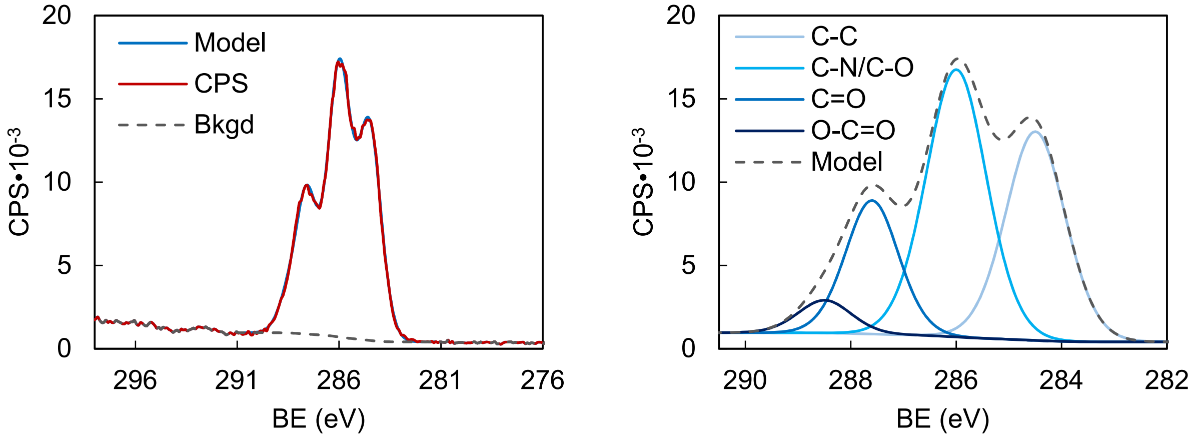


*50% βGal$^{TEA}$*

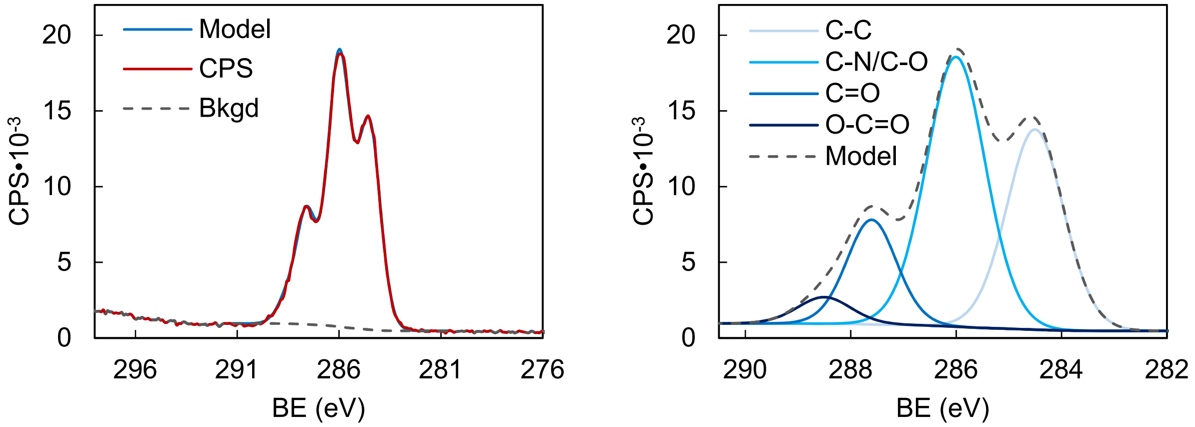


*50% α1,2Fuc-βGal*$^{TEA}$

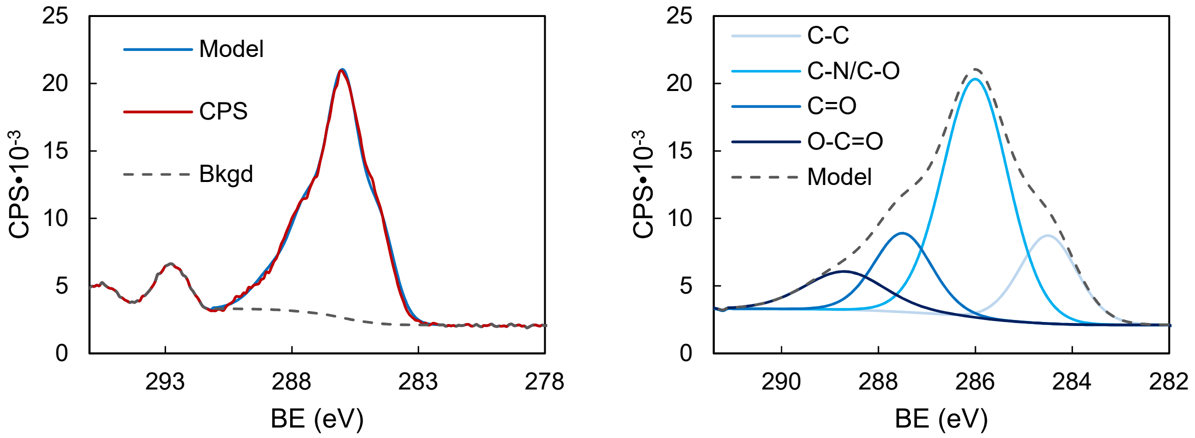


*50% βGal*$^{TEA}$

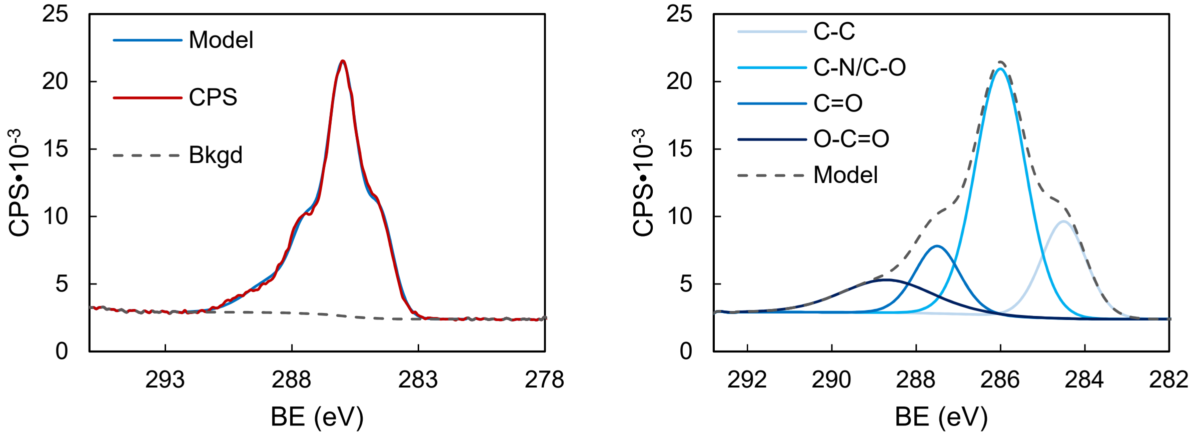


*50% α1,3Fuc-βGal*$^{TEA}$

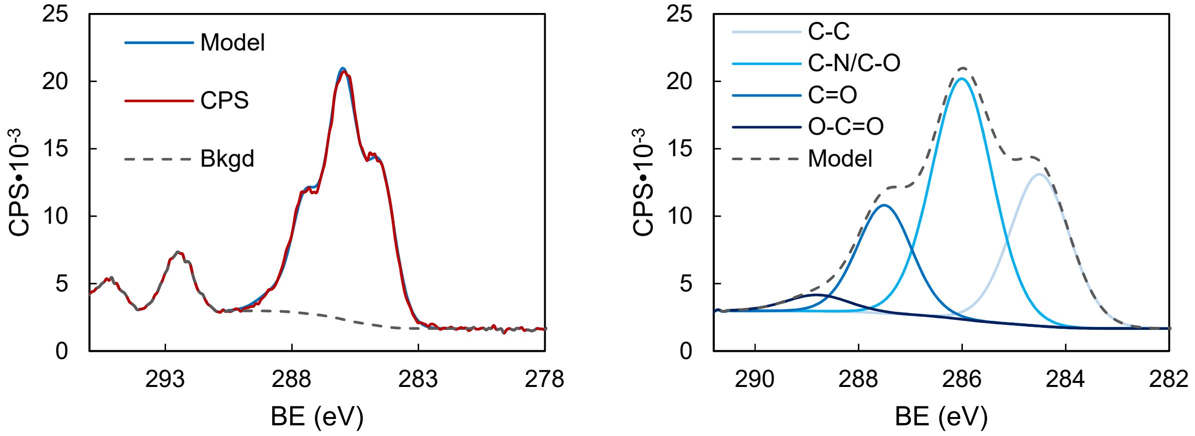


*50% βGal[TEA]*

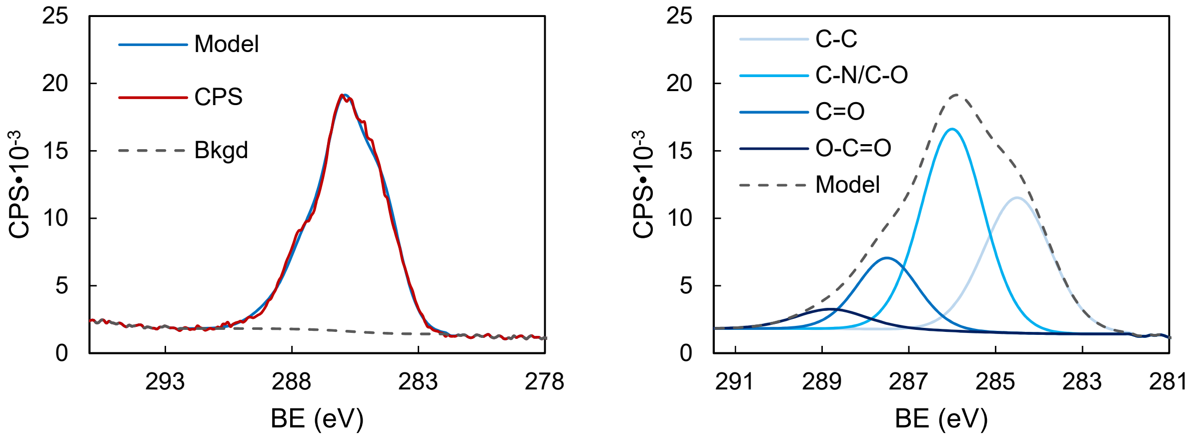


*50% α1,4Fuc-βGal[TEA]*

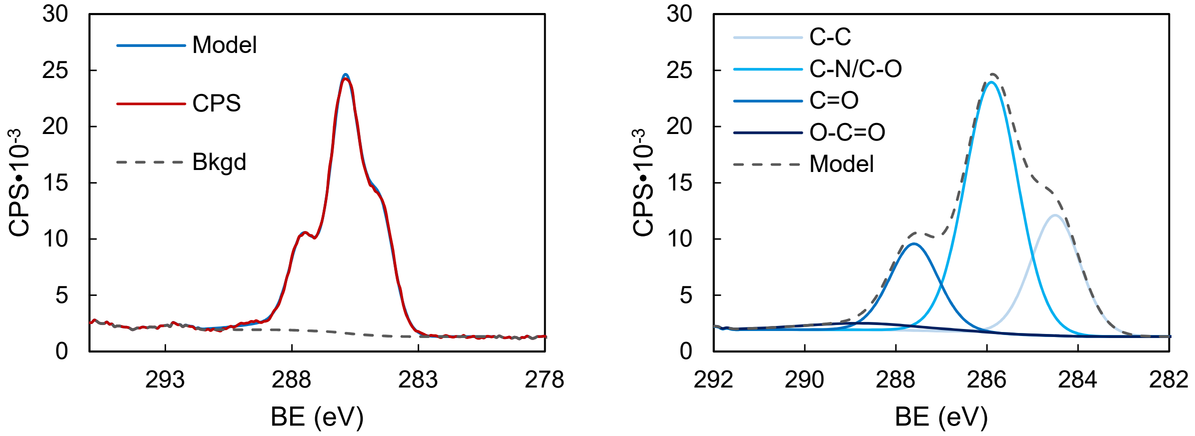


*50% βGal[TKA]*

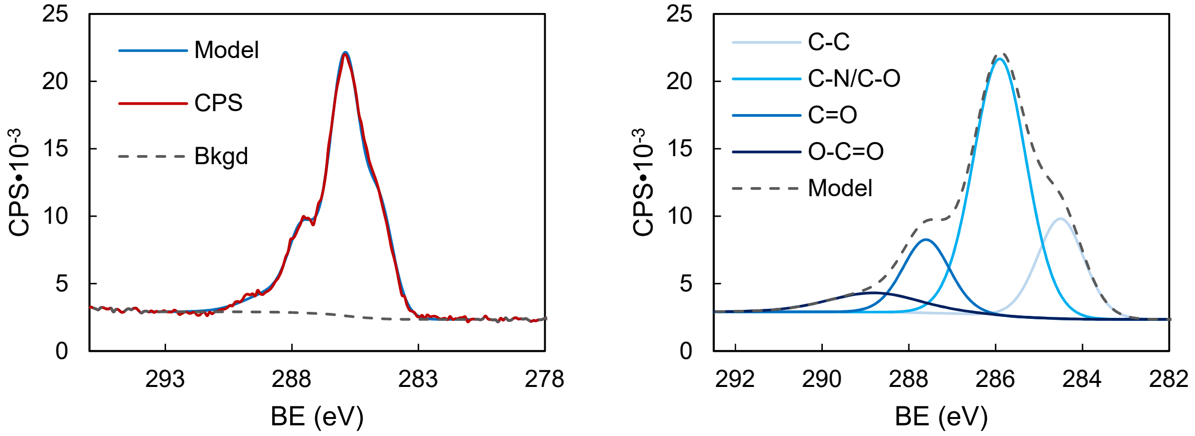


*50% α1,2Fuc-βGal$^{TKA}$*

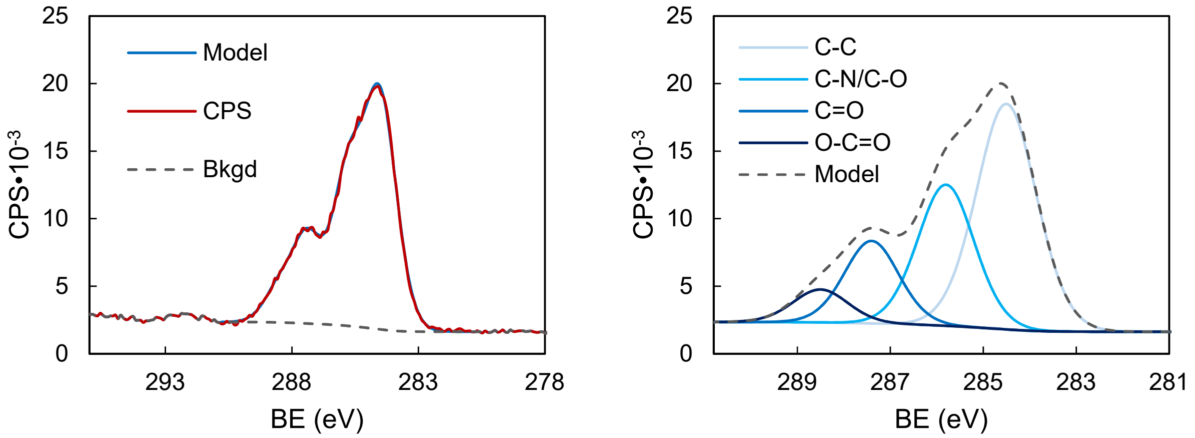


*25% βGal$^{TEAP}$*

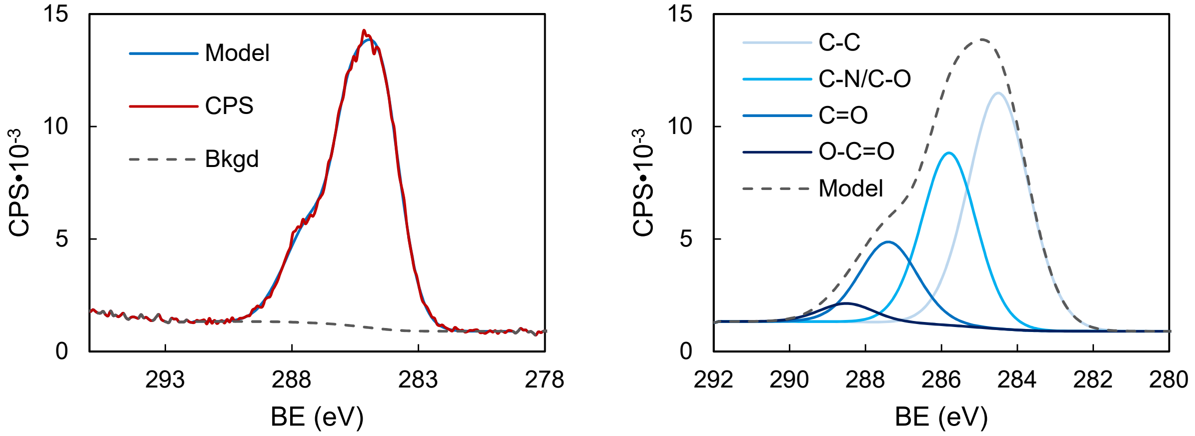


*25% α1,2Fuc-βGal$^{TEAP}$*

**Mucinase Digest Gels**

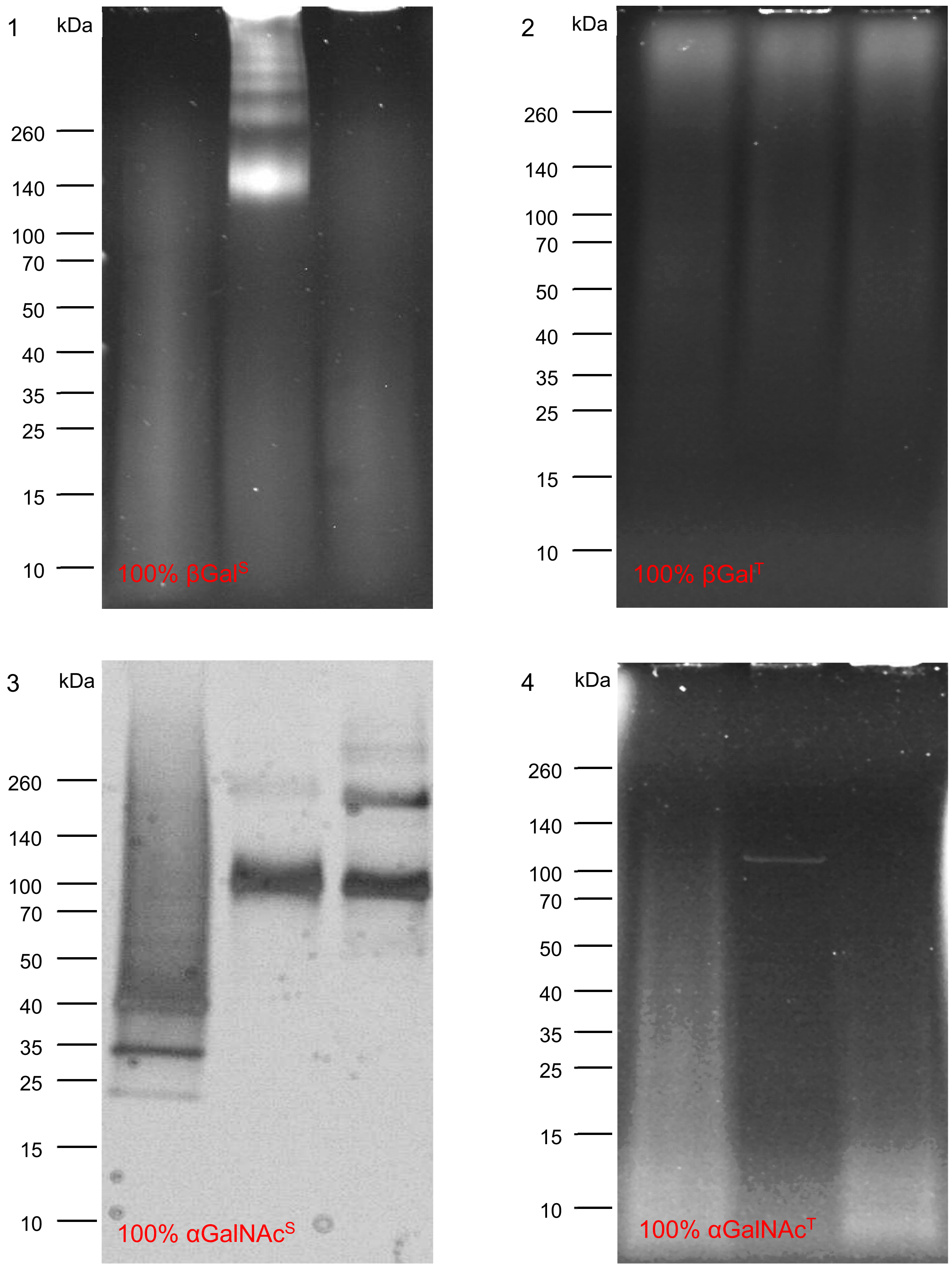

5

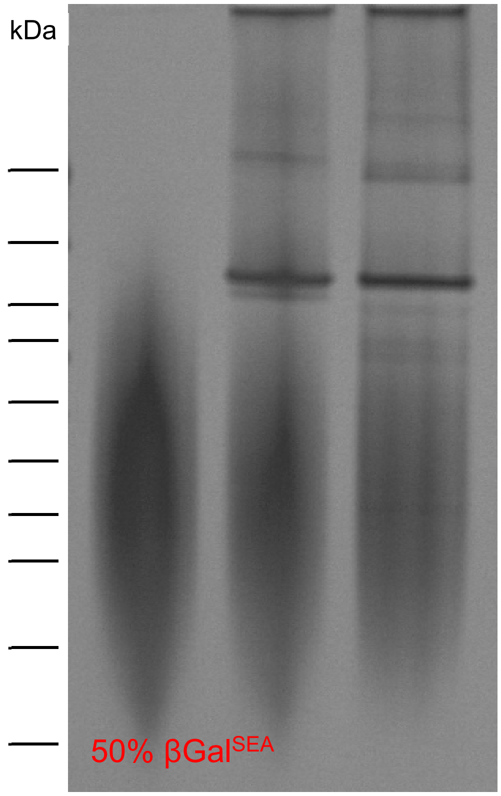
kDa
260
140
100
70
50
40
35
25
15
10
50% βGal^SEA

6

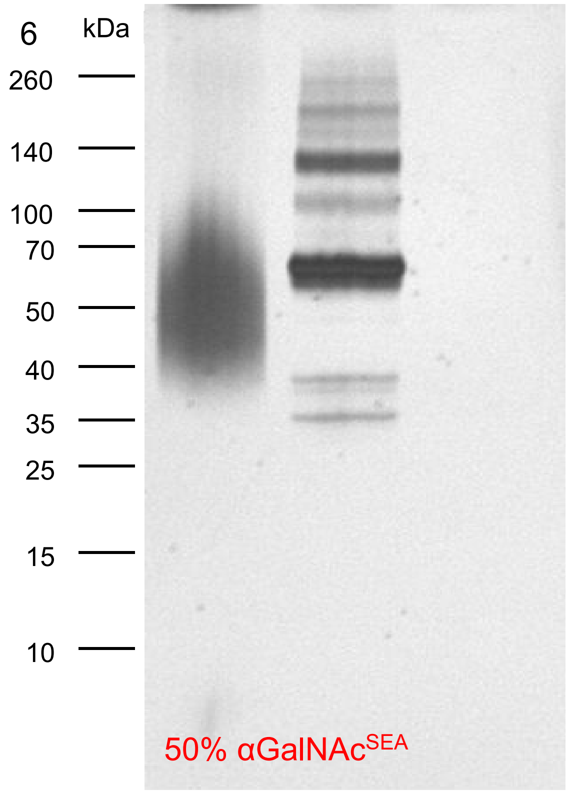
kDa
260
140
100
70
50
40
35
25
15
10
50% αGalNAc^SEA

7

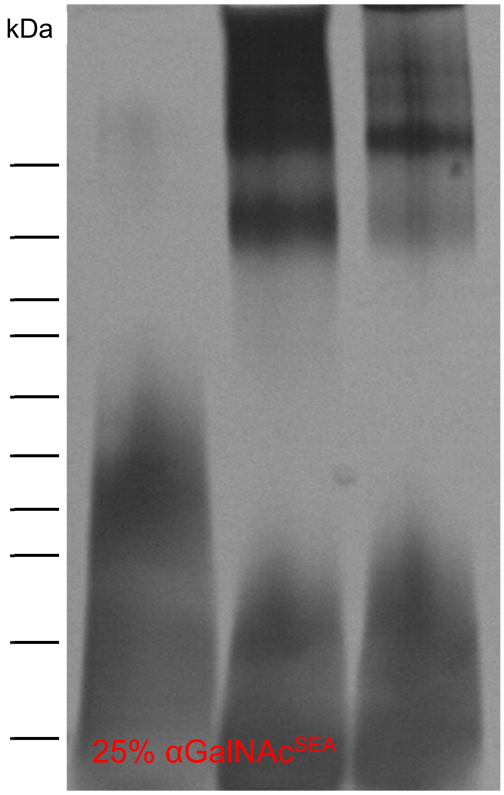
kDa
260
140
100
70
50
40
35
25
15
10
25% αGalNAc^SEA

8

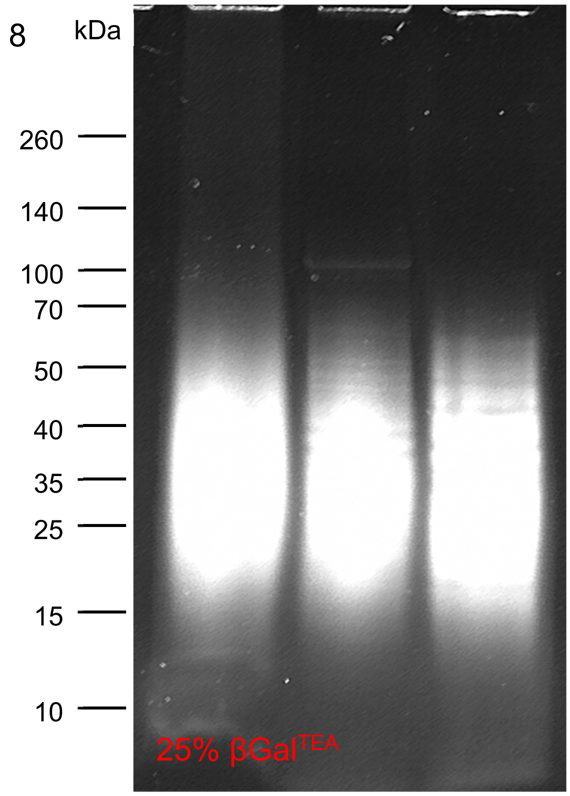
kDa
260
140
100
70
50
40
35
25
15
10
25% βGal^TEA

9

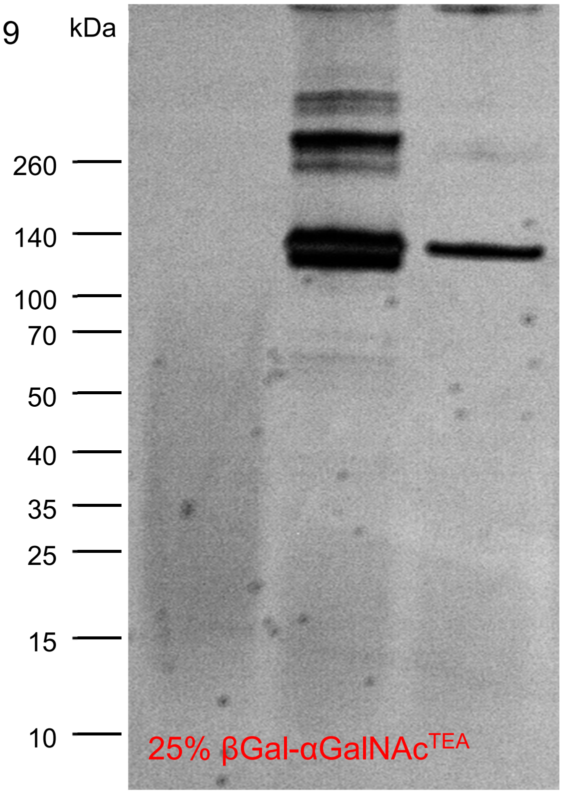
kDa
260
140
100
70
50
40
35
25
15
10
25% βGal-αGalNAcTEA

10

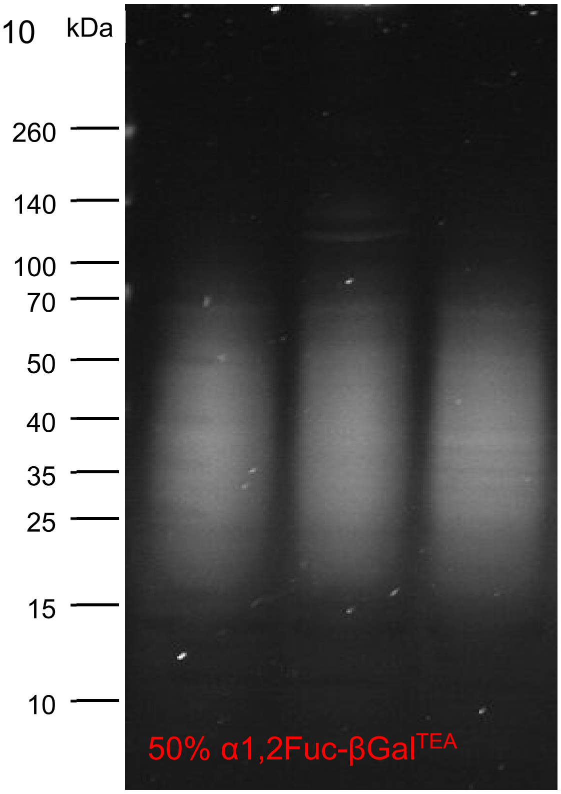
kDa
260
140
100
70
50
40
35
25
15
10
50% α1,2Fuc-βGalTEA

11

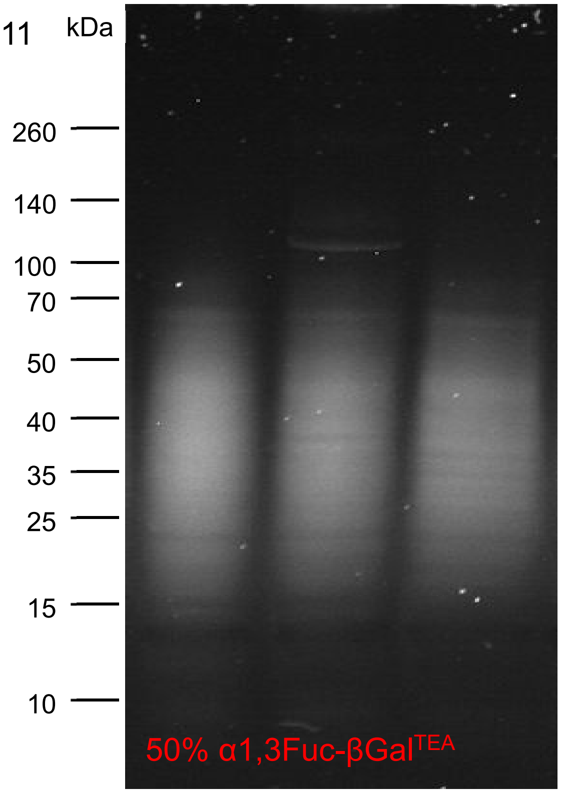
kDa
260
140
100
70
50
40
35
25
15
10
50% α1,3Fuc-βGalTEA

12

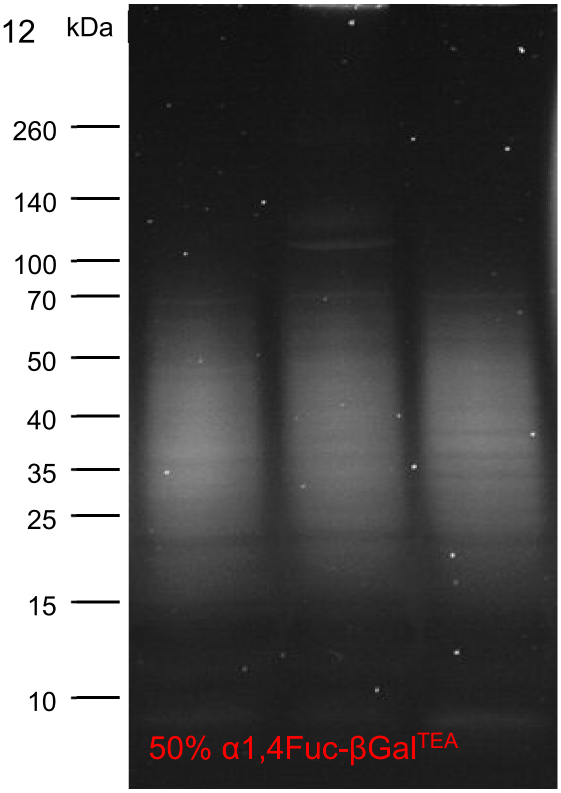
kDa
260
140
100
70
50
40
35
25
15
10
50% α1,4Fuc-βGalTEA

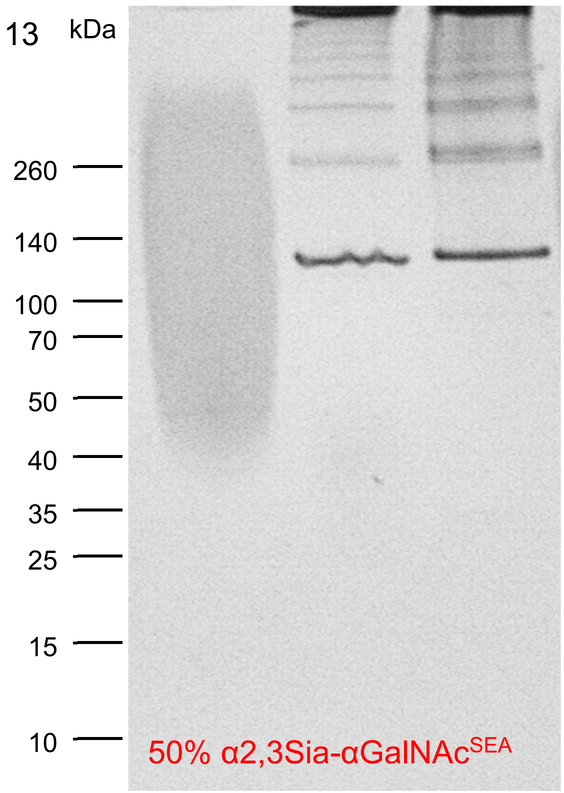
13
kDa
260
140
100
70
50
40
35
25
15
10
50% α2,3Sia-αGalNAc^SEA

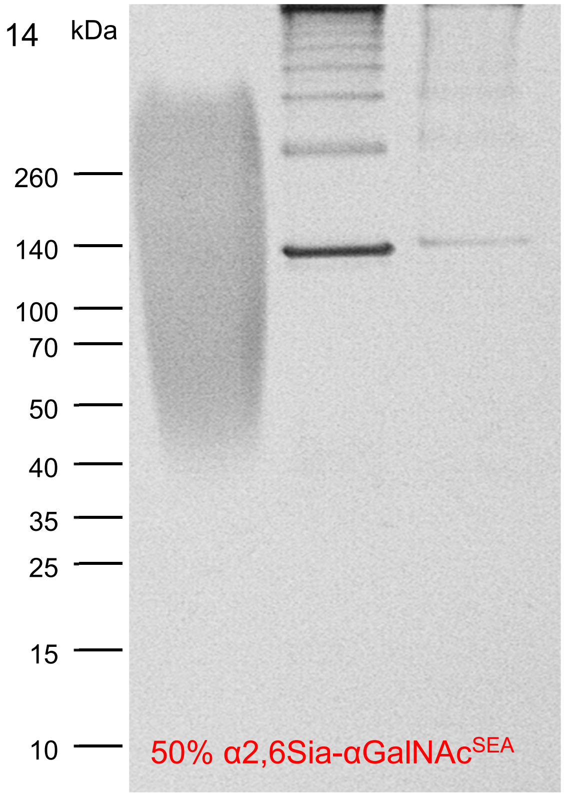
14
kDa
260
140
100
70
50
40
35
25
15
10
50% α2,6Sia-αGalNAc^SEA

(12) Yu, H.; Chokhawala, H. A.; Huang, S.; Chen, X. One-Pot Three-Enzyme Chemoenzymatic Approach to the Synthesis of Sialosides Containing Natural and

Non-Natural Functionalities. *Nat Protoc* **2006**, *1* (5), 2485–2492. https://doi.org/10.1038/nprot.2006.401.
